# Developing a general AI model for integrating diverse genomic modalities and comprehensive genomic knowledge

**DOI:** 10.1101/2025.05.08.652986

**Authors:** Zhenhao Zhang, Xinyu Bao, Linghua Jiang, Xin Luo, Yichun Wang, Annelise Comai, Joerg Waldhaus, Anders S. Hansen, Wenbo Li, Jie Liu

## Abstract

Advances in next-generation sequencing technologies have vastly expanded the availability of diverse genomic, epigenomic and transcriptomic data, presenting the opportunity to develop a general AI model that integrates comprehensive genomic knowledge into a unified model. Unlike previous predictive models, which are typically specialized to certain tasks, our general AI model unifies a wide range of genomic modalities, such as nascent RNA and ultra-high-resolution chromatin organization, within a multi-task architecture. Using ATAC-seq and DNA sequences as inputs, we incorporated diverse genomic modalities as output, and the model exhibits strong generalizability across different cell types and tissues in all tasks we trained. It accurately predicts gene-level transcription measured by various nascent RNA assays, and effectively captures enhancer-associated transcription. Additionally, it also accurately captures the potential functions of non-coding genetic variants and regulatory elements. Additionally, we extended the model trained on human data to a mouse general model, achieving accurate predictions of genomic modalities, such as high resolution chromatin contact maps with limited data availability, which are further validated using an established mouse inner-ear study. This comprehensive approach offers a powerful tool for understanding genome regulation in both human and mouse species.

## Introduction

Complementary next-generation sequencing assays have been developed to capture comprehensive facets of genomic activity. For instance, ATAC-seq measures chromatin accessibility, and ChIP-seq identifies histone modifications or the chromatin binding activities of transcription factors (TFs) and co-factors, whereas RNA-seq and GRO-seq assess RNA abundance and transcriptional activity [1, 2]. The complimentary data sources present a unique opportunity to develop computational models that integrate insights across diverse biological contexts to understand the complex, regulatory functions in the human genome. Recently, a number of computational models have been developed to predict multiple genomic modalities, either employing joint prediction strategies with a multi-task architecture for several modalities [3, 4, 5, 6] or utilizing pre-training and fine-tuning techniques to train various models for different downstream tasks [7, 8, 9, 10]. However, these multi-task predictive models typically gain information on a limited set of genomic tasks, and those pre-training models [8, 10] have individual models separately fine-tuned for different tasks. Consequently, these models struggle to integrate diverse genomic knowledge within a single unified framework. Therefore, a notable gap remains in the development of *a general artificial intelligence (AI) model* that concurrently accounts for different genomic modalities, embedding comprehensive genomic knowledge within a unified model. Exploring general models in the fields of computer vision and natural language processing [11, 12] indicates the potential for similar advancements in genomics.

To address this gap, we develop a general AI model that employs a multi-task architecture to jointly predict diverse genomic modalities. The general model embeds comprehensive genomic knowledge by leveraging rich datasets with a wide array of epigenomic features and experimental assays from a large number of cell lines and tissues, including comprehensive ChIP-seq datasets, and high-resolution 3D chromatin organization maps, as well as the direct outputs of the genome function, i.e., nascent and mature RNAs. Indeed, our model accurately predicts nascent RNA production at both the gene and enhancer levels, demonstrating its potential to make reliable predictions at the pseudobulk and tissue levels, where nascent RNA profiling protocols are often impractical. This aspect has not been explored by previous methods. Furthermore, our model has shown great improvement in characterizing non-coding regulatory elements, inferring their functions with high precision, which are validated through experimental studies such as eQTL mapping, lentiMPRA, and CRISPR perturbation [13, 14, 15].

In addition, we expanded the human general model to mouse. Indeed, an extensive array of genomic data is available across various species, particularly in mouse, which is an invaluable model to study human biology due to their close genetic and physiological parallels [16]. Despite this, a notable gap in the field is the absence of a model capable of predicting multiple genomic modalities in mouse. Recognizing the importance [17], we adapted our human-centric model and developed a general model for mouse species. Fine-tuning on mouse genomic data was necessary to accommodate unique characteristics in epigenomics and transcriptomics specific to mouse. The proposed mouse general model predicts critical regulatory chromatin interactions that are hardly detected in existing experimental chromatin contact maps, providing a valuable alternative to region-capture methods [18] that are constrained by the limited regions. The model effectively captures histone mark dynamics during developmental transitions and cell type-specific expression patterns in mouse tissues. A case study in the mouse inner ear further validates its utilities to capture biologically meaningful insights.

## Results

### A general model for multi-task learning across diverse genomic modalities to embed comprehensive and generalizable knowledge

We developed a general AI model that integrates a broad spectrum of genomic data in a multi-task architecture (Figure 1a,b). Compared with previous multi-task models [3, 4, 19], our general model predicts a larger, more comprehensive collection of genomic modalities simultaneously. It includes over 1,000 transcription factors (TFs), eleven commonly profiled histone marks, common mRNA assays (e.g., CAGE-seq and RNA-seq), nascent RNA assays (e.g., GRO-seq [2], Bru-seq [20] and TT-seq [21, 22]), and high-resolution 3D chromatin contact maps (e.g., Intact Hi-C [23] and Micro-C [24]), across a diverse set of cells and tissues (Figures 1f and S4a). Certain modalities are difficult to obtain due to specialized protocols (e.g., nascent RNA) or prohibitive costs (e.g., Micro-C). However, our general model can effectively impute these modalities when they are included in the training, enabling more comprehensive analyses in the situations where direct profiling is difficult.

**Fig. 1.**
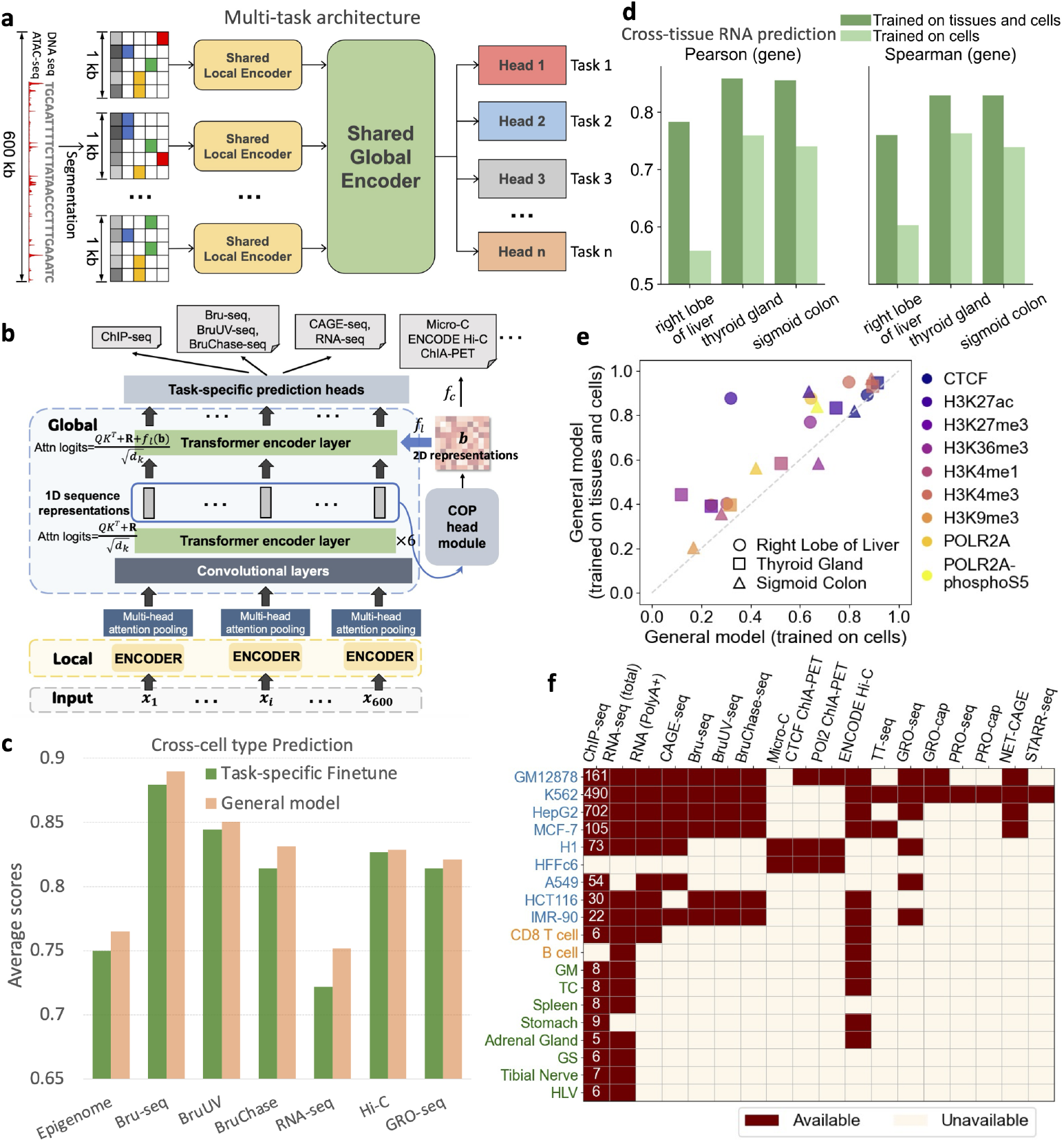
A general model to predict multiple genomic modalities from chromatin accessibility. **a**, The general model employs a multi-task architecture to simultaneously predict multiple genomic modalities. The input, consisting of a 600kb DNA sequence and ATAC-seq data, is segmented into 1kb bins. These bins are processed through a shared local encoder and a global encoder, followed by task-specific heads that predict various modalities at a 1kb resolution, with the exception of ChIA-PET predictions which are made at a 5kb resolution. **b**, The model integrates information across different modalities. It learns 1D representations through local encoders, which are then processed by convolutional layers and six transformer encoder layers with relative positional encoding in the global encoder. 2D features derived from these 1D representations are processed by a Chromatin Organization Prediction (COP) head module (illustrated in Figure S4e) to predict chromatin contact maps. These 2D features are also utilized in the final transformer encoder layer to refine the 1D sequence representations for predicting other modalities. **c**, The general model performs comparably to or better than task-specific fine-tuned models. For each modality used in the comparison, the two models use the same cell line data in training and testing (Figure S3a). Reported are average performance across different testing cell types on chromosomes 10 and 21. Spearman’s correlation is employed in the RNA-seq prediction task, whereas Pearson’s correlation is utilized in other tasks. **d**, The general model, trained on both cell lines and multiple tissue data, outperforms the general model trained only on cell lines in cross-tissue RNA-seq prediction. Pearson’s correlation and Spearman’s correlation are calculated across bins associated with genes on chromosomes 10 and 21. **e**, The general model, trained on both cell lines and multiple tissue data, outperforms the general model trained on cell lines in predicting cross-tissue epigenome features. Pearson’s correlation coefficients across the genome on on chromosomes 10 and 21 are reported. **f**, Overview of the modalities and the range of cell and tissue types incorporated to train the general model. Cell lines are colored in blue, primary cells are in orange, and tissues are in green. The numbers of ChIP-seq profiles used are indicated.

Our general model effectively learns general and universal sequence representations, distinguishing it from previous pre-training models that employ separate fine-tuned models for individual tasks [10, 8]. To assess the training effectiveness, we compared it with task-specific models trained based on the EPCOT framework [10], which are initially pre-trained and then fine-tuned for each downstream task using the same model architecture (i.e., the same number of parameters) and the same training data (Figure S2a). We focused on cross cell-type evaluation to test generalizability, and the evaluation spans a broad range of genomic modalities (Figure S3a). Across seven prediction tasks, our model either performed either comparably to or better than the task-specific models, with a significant improvement in RNA-seq prediction (Figure 1c).

Many of the existing predictive models are trained with cell line data. To capture biological insights beyond cell lines, we have incorporated multiple human tissue sequencing datasets available from EN-Tex [25] and Intact Hi-C contact maps from ENCODE [23] (Figure 1f). Tissues more closely resemble human’s complex biological processes compared to in vitro cell lines, and offer insights that are crucial for human diseases. We found that our current general model that incorporates multiple tissue data outperforms the model that was trained using only cell line data in epigenome and RNA-seq prediction tasks on unseen tissues (Figures 1d,e and Figures S3c,d). Furthermore, this general model also outperforms average experimental signals across multiple cells and tissues included in the training, which is a strong baseline in cross-cell type evaluation [26]. However, the improvement in Intact Hi-C prediction was less significant, though both models still surpassed the average cell and tissue Hi-C contact maps (Figure S3e,f). Further-more, we observed that integrating extra tissue data did not compromise the model’s generalizability to unseen cell lines (Figure S3g).

### The general model accurately predicts cell-type specific nascent RNA signals including enhancer RNAs

Transcription is the first layer of gene expression control and is the direct consequence of epigenomic and genomic regulation at the chromatin level [27]. Transcription can be accurately measured by nascent RNA sequencing assays such as GRO/PRO-seq, Bru-seq and TT-seq [27]. The transcriptional signals from non-coding regulatory regions such as enhancer RNAs (eRNAs) are also highly correlated with the functions and activity of such elements [28, 29]. However, most existing expression predictive models focus on mRNAs instead of nascent RNAs [3, 6, 30]. In contrast, our general model predicts nascent RNAs such as GRO-seq [2] and GRO-cap [31], which is a critical but currently missing prediction task. Furthermore, our model uses ATAC-seq data as inputs, enabling the prediction of multiple experimental assays that measure nascent RNA for unseen cells and tissues. In contrast, although several sequence-based models have been developed to predict GRO-cap or PRO-cap data [32, 33] from DNA sequences for transcription initiation interpretation, these models fail to impute data for new cells or tissues as they only use DNA sequences as input, and therefore are cell-type agnostic. Our model overcomes this limitation, making it essential to evaluate its effectiveness across different cellular contexts. To comprehensively assess its predictive capability, we evaluated nascent RNA predictions at *both gene and candidate enhancer levels*.

At the gene level, we tested how well it predicts Bru-seq data for five unseen cell lines. Notably, three of these lines (Caco2, Calu3, and A673) lack available ATAC-seq profiles in ENCODE, so we instead utilized pseudobulk snATAC-seq data from ENCODE as the input. Our model outperformed the baseline, defined as the average experimental signal from multiple training cell types (Figure 2a). Similarly, we evaluated how well it predicts strand-specific GRO-seq and TT-seq data using chromatin accessibility data (either bulk ATAC-seq or DNase-seq) from various studies [34, 35, 36]. This assessment tested the model’s robustness in generalizing across different chromatin accessibility data sources. Across GRO-seq and TT-seq experiments, the model consistently outperformed the average signal baseline, demonstrating superior predictive accuracy across genomic bins in gene regions, as validated on diverse datasets (Figure 2b,c). Additionally, we tested whether our trained model is sensitive enough to capture the transcriptional changes resulted from acute cellular signals. We applied our general model to predict transcriptional changes at nascent RNA levels in an unseen cell type MCF10A treated with a well established TGF*β*-1 signal. This signal is known to rapidly regulate gene transcription and change cell motility and migration [37]. We predicted TT-seq from ATAC-Seq data of MCF10A cells with and without 24 hours of TGF*β*-1 treatment [38]. We also experimentally conducted TT-seq in the same setting as a validation. Our model predicts nascent RNA well, achieving higher correlation scores than the average signal baseline (Figure 2c). The model also effectively captured transcriptional changes induced by this acute signal treatment (see one example region in Figure 2d). We further compared the global transcriptional changes predicted by our model versus experiment. Using log-transformed fold changes between the 0-hour and 24-hour TGF*β*-1 treatments across differentially expressed genes, we observed an overall satisfactory correlation between our model predictions and experimental results (Figure S6). However, some gene-specific changes were not well predicted (Figure S6), which could be attributed to the fact that the ATAC-seq data we used for prediction and the experimental TT-seq data are derived from separate labs.

**Fig. 2.**
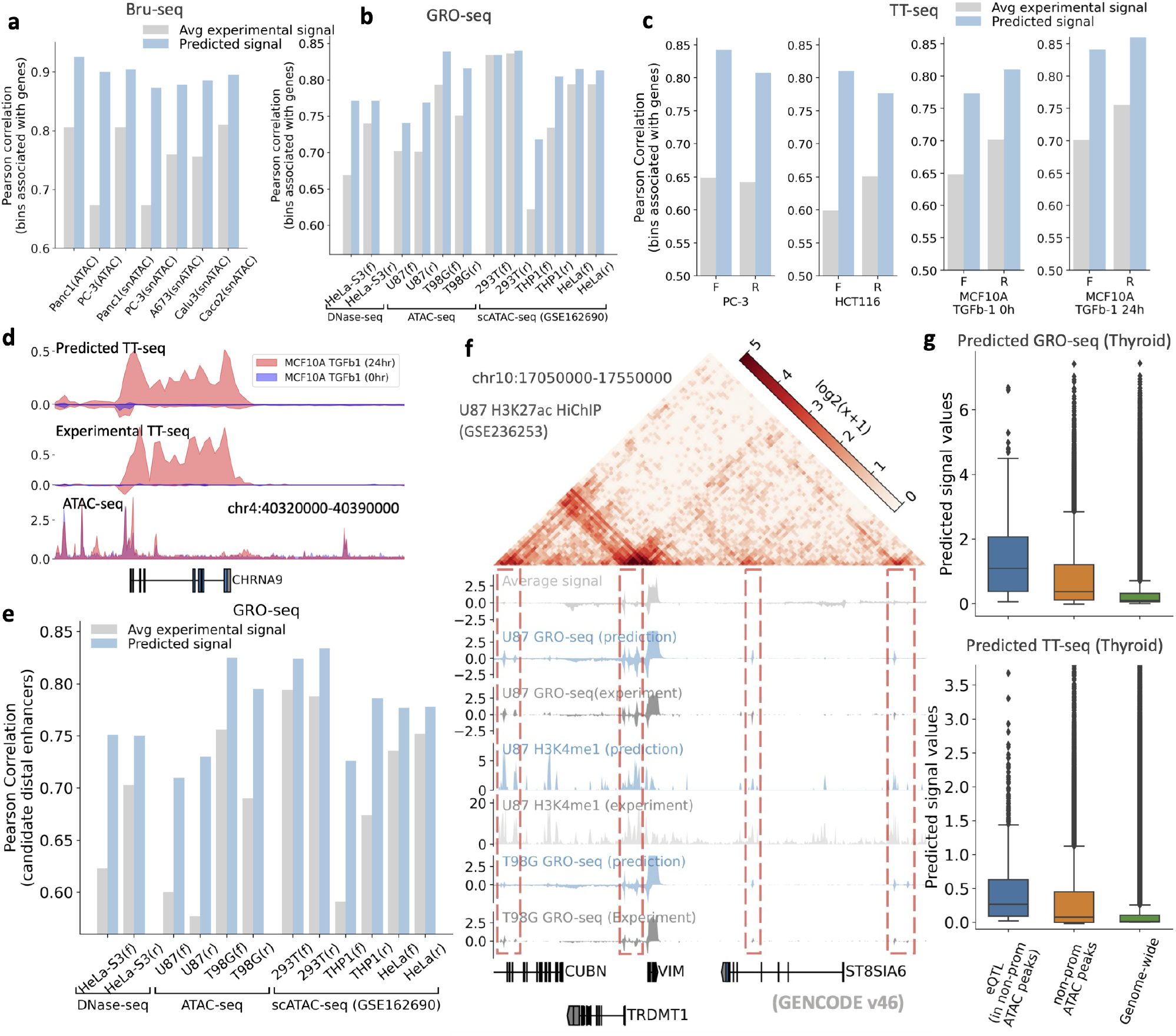
General model accurately predicts nascent RNA at the gene level and across enhancer regions. The general model outperforms the average signal from experimental data across genomic bins associated with gene regions for **a**, Bru-seq predictions, using snATAC-seq and ATAC-seq in PC-3 and Panc1 and snATAC-seq alone for the other three cell lines; **b**, strand-specific GRO-seq signals, leveraging DNase-seq and ATAC-seq; and **c**, strand-specific TT-seq in two cell types and MCF10A cells with and without treatment. **d**, The model accurately predicts the expression changes in the CHRNA9 gene between 0-hour and 24-hour TGF*β*-1 treatments, effectively capturing subtle changes in the input ATAC-seq data. **e**, Predicted GRO-seq signals outperform average experimental signals across candidate distal enhancer regions, using DNase-seq, ATAC-seq, or pseudobulk scATAC-seq data. **f**, A specific region demonstrates that the general model accurately predicts putative eRNAs (highlighted in red dashed boxes) in two GBM cell lines. These are not well detected by the average experimental signal. The U87 H3K27ac HiChIP data further validates that these predicted signals represent eRNAs. **g**, Tissue-level predictions are validated using GTEx eQTLs. eQTLs with a PIP score greater than 0.1 that overlap with corresponding ATAC-seq peaks are utilized. Predicted GRO-seq and TT-seq signal values in thyroid tissue are visualized in a box plot for eQTL bins that do not overlap with promoter regions, along with all non-promoter ATAC-seq peak bins. Additionally, signal values across all genomic bins are presented for background comparison (labeled as “Genome-wide”). The general model consistently predicts higher signal intensities at eQTL loci, including those in putative enhancer regions.

At enhancer regions, we evaluated eRNA predictions across bulk, psuedo-bulk (single-cell clusters), and tissue levels, respectively. At the bulk level, across GRO-seq, Bru-seq, BruUV-seq, TT-seq, and NET-CAGE datasets, our model outperformed the baseline — average signal across candidate enhancer regions (Figures 2e, S7, S8). By incorporating high-resolution chromatin contact maps (H3K27ac HiChIP and Hi-C), the model also showed potential in linking transcribed enhancer regions to neighboring genes (Figures 2f and S5). For single-cell cluster-level predictions, we used pseudobulk ATAC-seq data from scATAC-seq for three cell types (293T, THP1, and HeLa) [39], with corresponding bulk GRO-seq data for validation. The model consistently outperformed the average bulk signal in predicting GRO-seq across genes and candidate distal enhancer regions (Figure 2e), demonstrating its ability to capture eRNA activity in the single-cell clusters. At the tissue level, due to limited nascent RNA sequencing data, we validated eRNA predictions using eQTLs, which are enriched in enhancer elements [40, 41], along with ATAC-seq data to identify open chromatin regions while excluding promoter-overlapping regions, thus focusing on putative enhancer regions. This approach leverages the well-established association between eRNA regions and eQTLs [42, 43], highlighting that genetically important enhancers (marked by eQTLs) are more frequently associated with active transcription. We hypothesized that, if an eQTL overlaps a non-promoter open chromatin region, it is likely located in a putative active enhancer, and thus our model would predict stronger nascent RNA (eRNA) signals at these eQTL loci compared to randomly selected non-promoter open regions and the genome-wide background. To test this, we gathered eQTL data from the eQTL Catalogue [13] and predicted GRO-seq, TT-seq, and NET-CAGE signals in five unseen GTEx tissues. Predictions were evaluated across three region types, (1) eQTL loci specifically within non-promoter open chromatin regions, (2) all non-promoter open chromatin regions, and (3) the entire genome. In these comparisons, (1) highlights putative active non-coding regions, while (2) and (3) serve as baselines for putative enhancer regions and the entire genome, respectively. The model consistently predicted significantly stronger nascent RNA signals at eQTL loci than at the baseline regions across all tissues (Figures 2g, S9). Notably, the model predicted statistically stronger signals at the putative active enhancer regions in (1) compared to the broader putative enhancer regions in (2), with support from *t*-test *p*-values (Figure S10). These findings highlight the model’s potential as a powerful tool for predicting eRNAs across cell types, single-cell clusters, and tissues when ATAC-seq data is available.

### The general model characterizes functions of non-coding regions in cell-type specific contexts

Non-coding regions of the genome harbor various regulatory elements that are crucial in gene regulation. Although multiple computational methods [3, 19, 14, 6] have been proposed to study the functions of non-coding elements, most of these methods are sequence-based models that rely solely on DNA sequence and derive genomic knowledge from a limited number of genomic modalities. Here, we explore the general model’s potential in studying variants and elements with datasets that were not used in model training, including eQTLs in putative non-coding regions, lentiMPRA data, and CRISPR perturbation data.

We first applied our general model to assess whether non-coding variants functionally regulate target genes in specific cell or tissue contexts. Given that eQTLs overlapping open chromatin regions are more likely to be functional regulatory variants, we evaluated our model’s ability to distinguish these functional eQTLs from other non-coding variants. To achieve this, we leveraged eQTL datasets in eQTL Catalogue [13], identifying putative non-coding variants by intersecting those variants with ATAC-seq peaks. We then categorized these variants into positive (functional eQTLs) and negative ones based on PIP values from SuSiE’s fine-mapping procedure, following a similar approach to prior models Enformer [3] and Borzoi [6]. Unlike sequence-based models such as Enformer and Borzoi, our general model integrates cell-type-specific bulk ATAC-seq or pseudo-bulk scATAC-seq data, allowing us to evaluate eQTL classification performance in novel cell contexts. Specifically, we trained our model on GTEx datasets [40] and tested it on 13 new cell types from five independent studies [44, 45, 46, 47, 48], each with available ATAC-seq data (Figure S12b,c). For this evaluation task, we developed a classifier utilizing the features of the general model and additional features such as Borzoi variant scores, to predict the regulation relationship between gene-variant pairs (Figure 3a, see Methods for details). We used Enformer and Borzoi which applied a random forest classifier on variant scores as baselines and further enhanced their variant scoring with a distance-to-gene TSS feature. Our model consistently outperformed these baselines in cross cell-type eQTL predictions, and also demonstrated superior performance on distal non-coding variants (Figure 3b), indicating its ability to predict non-coding functional eQTLs in unseen cell contexts. We further evaluated whether the informative representations learned by our general model could enhance eQTL classification in specific tissues. Following a similar approach in Enformer and Borzoi, we utilized our sequence representation of the variant bin and the 2D representation of the interaction between the variant and gene TSS bins, along with Borzoi’s variant scores, to train a random forest classifier to discriminate functional eQTLs for each tissue. We observed that integrating the learned general representations outperforms both Borzoi and Enformer in terms of ROC and PR curves across five unseen tissues (Figure S12e).

**Fig. 3.**
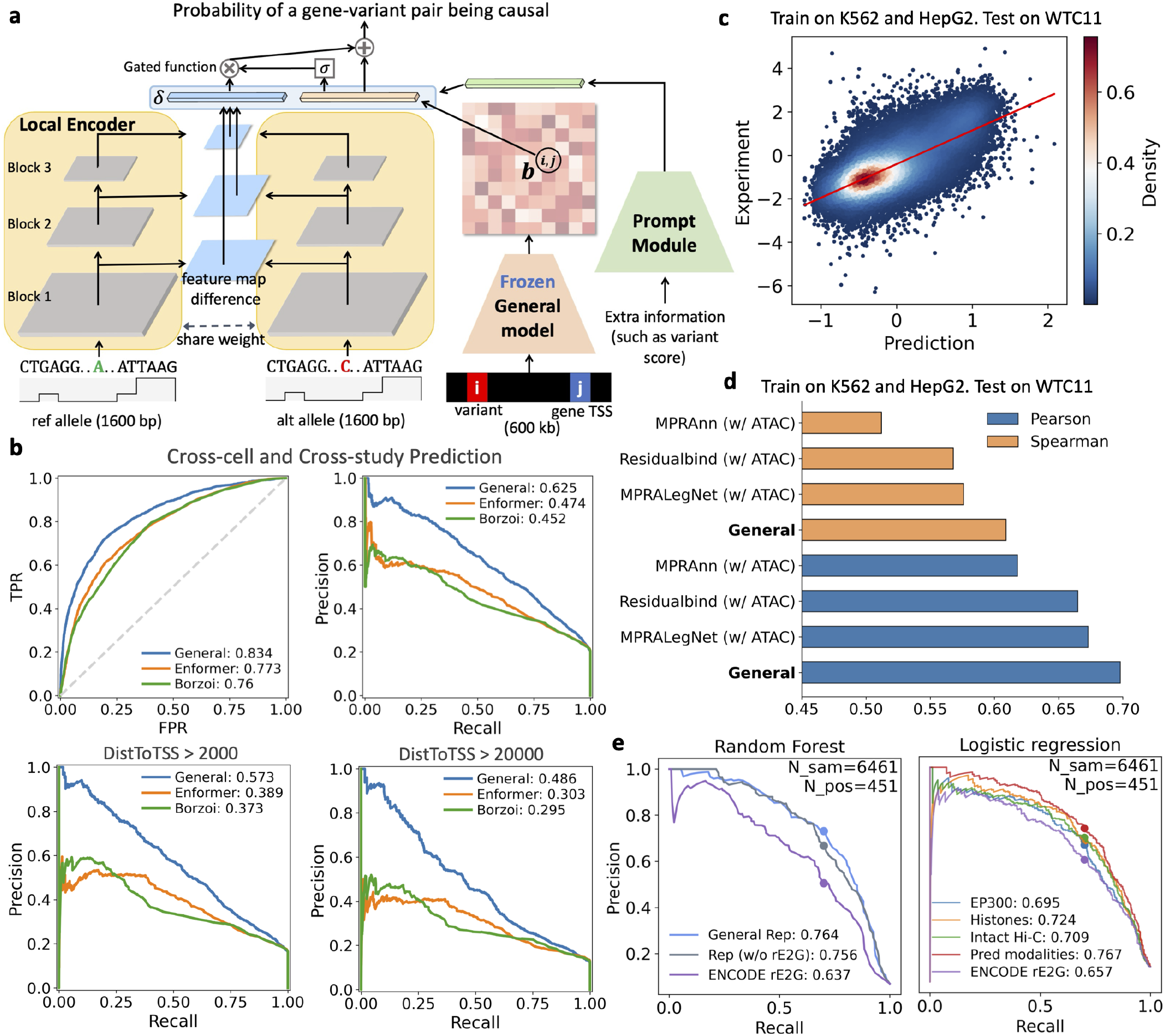
The general model elucidates the functions of non-coding variants and regulatory regions. **a**, The model architecture of the eQTL classification task. The local encoder of the general model is used to capture the local feature difference *δ* between reference and alternative alleles, which is then integrated with 2D interaction feature *b*_*i,j*_ between the variant bin *i* and gene TSS bin *j*, along with prompt features such as variant scores to predict the probability of a gene-variant pair being casual. The architecture of the prompt module is illustrated in Figure S12a. **b**, The general model more accurately predicts eQTLs in unseen cell types from various studies than both Enformer and Borzoi. **c**, The general model precisely predicts cross-cell type lentiMPRA activity scores. We present both predicted and experimental regulatory scores in WTC11, with the model trained on K562 and HepG2 cell lines. **d**, The general model surpasses MPRAnn, Residualbind, and MPRALegNet models in predicting cross-cell type lentiMPRA. **e**, The representations and predicted genomic modalities improves ENCODE-rE2G’s performance on K562 perturbation data. Random forest is applied to the general model’s representations with and without incorporating 13 features from the ENCODE rE2G model. Additionally, Logistic Regression is applied to predicted features combined with ENCODE rE2G features. Precision scores at 70% recall are presented. ‘Pred modalities’ refers to multiple predicted features, including EP300, histone marks, CAGE-seq, RNA-seq, Bru-seq, BruUV-seq, GRO-seq, GRO-cap, NET-CAGE, Micro-C, and Intact Hi-C.

Additionally, we evaluated the model’s utility in predicting the regulatory functions of non-coding elements. We leveraged a lentiMPRA study, which measured the regulatory activity of multiple elements in K562, HepG2, and WTC11 cells [14]. Given that only short sequences were involved in the lentiMPRA study, we fine-tuned only the local encoder of the general model, adding convolutional and linear layers to predict regulatory scores on both strands of elements (Figure S11). For comparison, we included two end-to-end deep learning models, MPRAnn and MPRALegNet, from the LentiMPRA study [14], as well as a strong baseline model, ResidualBind [49]. All three models can be adapted to integrate ATAC-seq data for cross-cell type predictions by slightly modifying their architectures, enabling predictions for new cell types. To assess the model’s ability to predict regulatory scores of elements in a new cell type, we trained the models on K562 and HepG2, two of the training cell lines used in the general model, and subsequently tested them on WTC11. The general model consistently outperformed these baselines in cross-cell type predictions (Figure 3c,d), indicating the model’s capability of measuring cell-type specific regulatory activities.

Furthermore, we validated the effectiveness of our general model in identifying enhancer-gene regulatory pairs, using CRISPR perturbation data on K562 [15] and comparing it with the state-of-the-art method, ENCODE-rE2G [15]. ENCODE-rE2G [15] employs logistic regression with 13 features to identify regulatory elements, of which only one is a DNase-seq feature requiring cell-type-specific experimental measurements. Our model, by comparison, only requires ATAC-seq data as input, making it conceptually similar to ENCODE-rE2G and prompting us to investigate its potential for improvement. Given that K562 was initially a training cell line for the general model, we retrained the general model excluding K562, instead using data from seven other cells and two tissues (Figure S13a,d). To evaluate the utility of the model’s learned representations in identifying enhancer-gene interactions, we applied a random forest classifier to these representations. This approach yielded significant improvement over ENCODE-rE2G, both with and without incorporating its features (Figure 3e left panel), indicating that our informative representations are effective in detecting regulatory interactions. In addition, we assessed whether integrating predicted modalities with ENCODE-rE2G features could enhance the performance. We found that including different predicted modalities indeed provided a more accurate prediction of regulatory interactions, and integrating multiple predicted features yielded the highest AUPR scores (Figure 3e right panel). Furthermore, we introduced individual predicted TFs to the ENCODE-rE2G, with TFs like CUX1 showing the most significant performance improvement, supported by literature confirming its association with enhancer activity in K562 [50, 51] (Figure S13b). Additionally, these predicted features and learned representations could further enhance the performance of ENCODE-rE2G-extend which incorporates experimental histone marks, TFs, Hi-C contact maps, and other features (Figure S13c). This further demonstrates the utility of our general model.

### Transferring the general human model to a mouse model which accurately predicts multiple genomic modalities

Biomedical research frequently utilizes mouse models to understand human biology due to the high similarities between the two species in the genome [17], which indicates promise in transferring the human general model to mice. However, despite these similarities, crucial epigenetic and regulatory differences exist that limit the direct application of the human model to mouse studies [17, 52, 53]. To address this, we propose developing a mouse-specific general model adapted from the human model, providing a robust foundation for mouse studies. To our knowledge, no existing computational model offers this capability.

To achieve this, we adapted and fine-tuned the human model to create a general model for mice, incorporating a diverse array of mouse experimental data from various cells and tissues into training, including ChIP-seq, RNA-seq, CAGE-seq, PRO-seq, GRO-seq, NET-CAGE, Hi-C, and Micro-C chromatin contact maps (Figure S14 and Methods section titled ‘Training Mouse General Model’). In addition, we also included region capture Micro-C (RCMC) data [18], which offers significantly more detailed chromatin contact information than genome-wide Micro-C contact maps but covers only about 5Mb across five loci. The mouse general model achieved high accuracy in predicting these diverse genomic modalities.

We first evaluated its ability in predicting high-resolution RCMC data. Given the limited data availability, we established two baselines, (1) fine-tuning the entire human general model on the RCMC data, and (2) using linear probing to train a linear prediction layer on the fixed human model — an effective method in few-shot learning scenarios [54]. We used RCMC from a 420kb region on chromosome 6 for testing after training the model with four other loci. In the hold-out testing region, the general model outperformed both fine-tuning and linear probing in terms of Pearson and Spearman correlations (Figure 4a,b). Nevertheless, the predicted contact maps from all three methods revealed finer-resolution chromatin contacts than those observed with Micro-C data in this region (Figure S16a).

**Fig. 4.**
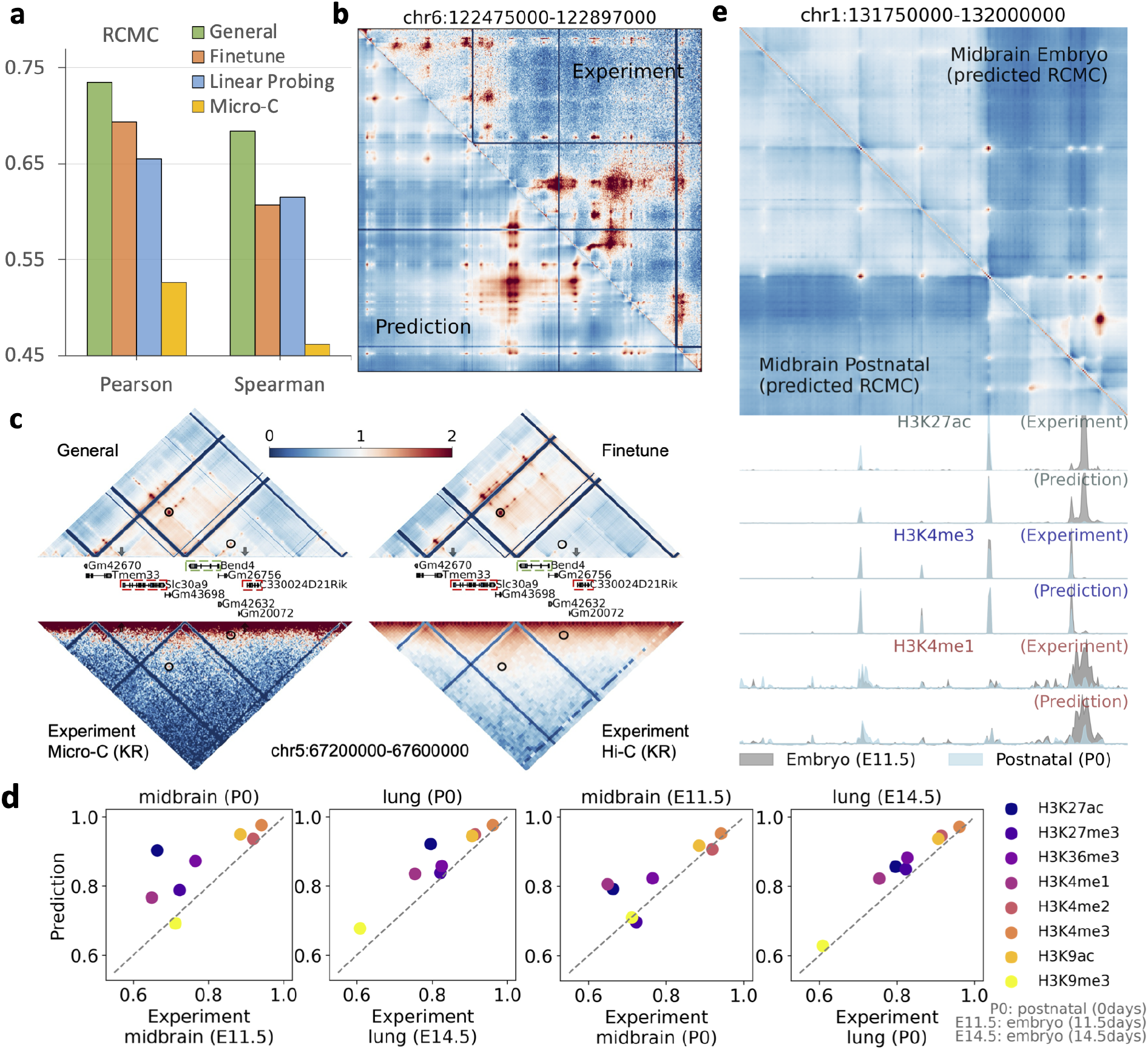
The mouse general model accurately predicts different genomic modalities. **a**, The mouse general model outperforms both fine-tuning and linear probing applied to a human pre-trained model in predicting RCMC. **b**, The general model effectively recovers RCMC within the test region. Visualizations of predictions from other approaches are provided in Figure S16a. **c**, The general model identifies two critical interactions (highlighted in black circles), validated by perturbation studies, that are not clearly delineated by the fine-tuned model, nor by experimental Micro-C and Hi-C data. **d**, The model captures histone mark variations across different developmental stages, predicting histones more accurately than using experimental data from a varying developmental stage. Pearson correlation scores are reported. **e**, An illustrative example where the general model detects changes in histone marks across different developmental stages, which are also reflected in the predicted RCMC contact maps.

To further validate the predicted RCMC contact maps, we utilized a study [55] that investigated the effects of knocking out promoters of lncRNA and mRNA on neighboring gene expression. This study identified four regulatory interactions within our prediction range that were undetected by Hi-C contact maps [56]. Our general model detected clear loop patterns for all four interactions, whereas the fine-tuned model detected three interactions, missing the interaction between Bend4 and C330024D21Rik (Figures 4c and S16b). Although linear probing also detected these interactions, the loop patterns were not as obvious as those from the general model. Furthermore, we checked whether the experimental Micro-C [57] could capture these interactions. We found that these interactions showed weak signals in the KR-normalized Micro-C contact maps, and they were not discernible as loop patterns, although they are visible in the observed-over-expected (O/E) ratio counts. These results underscore the effectiveness of our general model in using limited regions of RCMC to impute other regions. Additionally, the O/E normalized mESC Hi-C contact map [58] failed to detect all of them.

Furthermore, the mouse general model accurately predicts histone marks. We observed that the model can identify histone mark changes across different developmental stages in tissues (Figure 4e). During training, we incorporated experimental assays from both postnatal and embryonic stages of heart and liver tissues (Figure S14). For evaluation, we leveraged data from both postnatal stages and early embryonic time points in the midbrain and lung, as well as from the embryonic forebrain. We established a baseline by predicting histone marks using the experiment data in different developmental stages within the same tissue, for instance, using histone data from postnatal day 0 in lungs to predict embryonic (14.5 days) lung histone marks, and compared these predictions with those generated by the general model from corresponding ATAC-seq data. The general model showed superior performance genome-wide, and also in ENCODE candidate promoter and distal enhancer regions (Figures 4d and S15a,b). Additionally, we observed that the general model predicted cell-type specific marks like H3K27ac significantly better, especially in candidate distal enhancer regions. Furthermore, we validated the histone mark prediction performance by comparing the general model’s performance against average histone signals derived from all training cells and tissues (Figure S14). This comparison included tests on those testing tissues and one new cell line, G1E, where the general model consistently demonstrated strong Pearson’s correlation (Figure S15d). Since human models have been directly used for mouse histone mark prediction [10], we further assessed the efficacy of species-specific adaptations by comparing the predicted histone marks from the mouse and human general models (Figure S15c,d). The mouse-specific model significantly outperformed the human model. Considering the distribution differences between human and mouse experimental data, we retrained the final linear layer of the human model for prediction. Despite this adjustment, the mouse model still outperformed the human model, indicating the necessity of fine-tuning to learn mouse-specific knowledge (Figures S15d).

Additionally, we evaluated the model’s prediction performance on expression data by leveraging PolyA+ RNA-seq from midbrain and lung tissues in embryonic and postnatal cells. To evaluate, for a given developmental stage (e.g., embryo), we used experimental data from a different stage (e.g., postnatal) as a baseline. Predictions based on data from the same stage exhibited higher correlation scores, indicating that the model accurately predicts the tissue-specific expression patterns (Figure S17a). In addition, we sought to evaluate whether our model reliably predicts enhancer RNAs typically captured by nascent RNA assays such as PRO-seq. Because PRO-seq data are limited in mouse species, we used H3K27ac, a well-established active enhancer mark, as a surrogate for nascent transcription to evaluate our predictions. Specifically, we assessed the correlation between predicted PRO-seq signals and experimental H3K27ac levels across candidate enhancer regions. We observed that regions exhibiting high H3K27ac also displayed strong predicted PRO-seq signals, indicating our model’s capability to capture enhancer-associated transcription. These findings highlight the model’s potential for accurately capturing enhancer RNAs (Figure S17b).

### Elucidating cell type-specific epigenomic regulation in a mouse inner ear study

In order to examine whether the model allows us to explore cell type-specific transcriptional regulation *in vivo*, we applied the mouse general model to the datasets generated from a mouse inner ear study [59]. The study investigates the regulatory landscape of auditory hair cells and generated two age-matched datasets (one scATAC-seq and one scRNA-seq) from postnatal day 2 mouse cochlear ducts (Table S1). As the first task, we aggregated scATAC-seq cells into six pseudo-bulk profiles based on cell type annotation, and then assessed whether the predicted RNA profiles from the general model could sufficiently distinguish cell type-specific expression patterns. We observed that, for each experimental cell type (columns in Figure 5a), the highest correlation was observed with the predicted expression of the corresponding scATAC-seq cell type. Next, we checked the characteristics of each cell type’s marker genes. We computed the correlation between the predicted gene expression and the experimentally measured single-cell gene expression. For each cell type, we selected the top 200 genes with the highest correlation and performed a Gene Ontology (GO) biological process analysis using Gene Ontology [60] (Table S2). It was observed that the marker genes accurately captured the functions of each cell type. For instance, in hair cells, processes such as ‘neurofilament bundle assembly’ and ‘vestibular receptor cell differentiation’ exhibit high fold enrichment and low false discovery rate, suggesting that these processes characterize well hair cells.

**Fig. 5.**
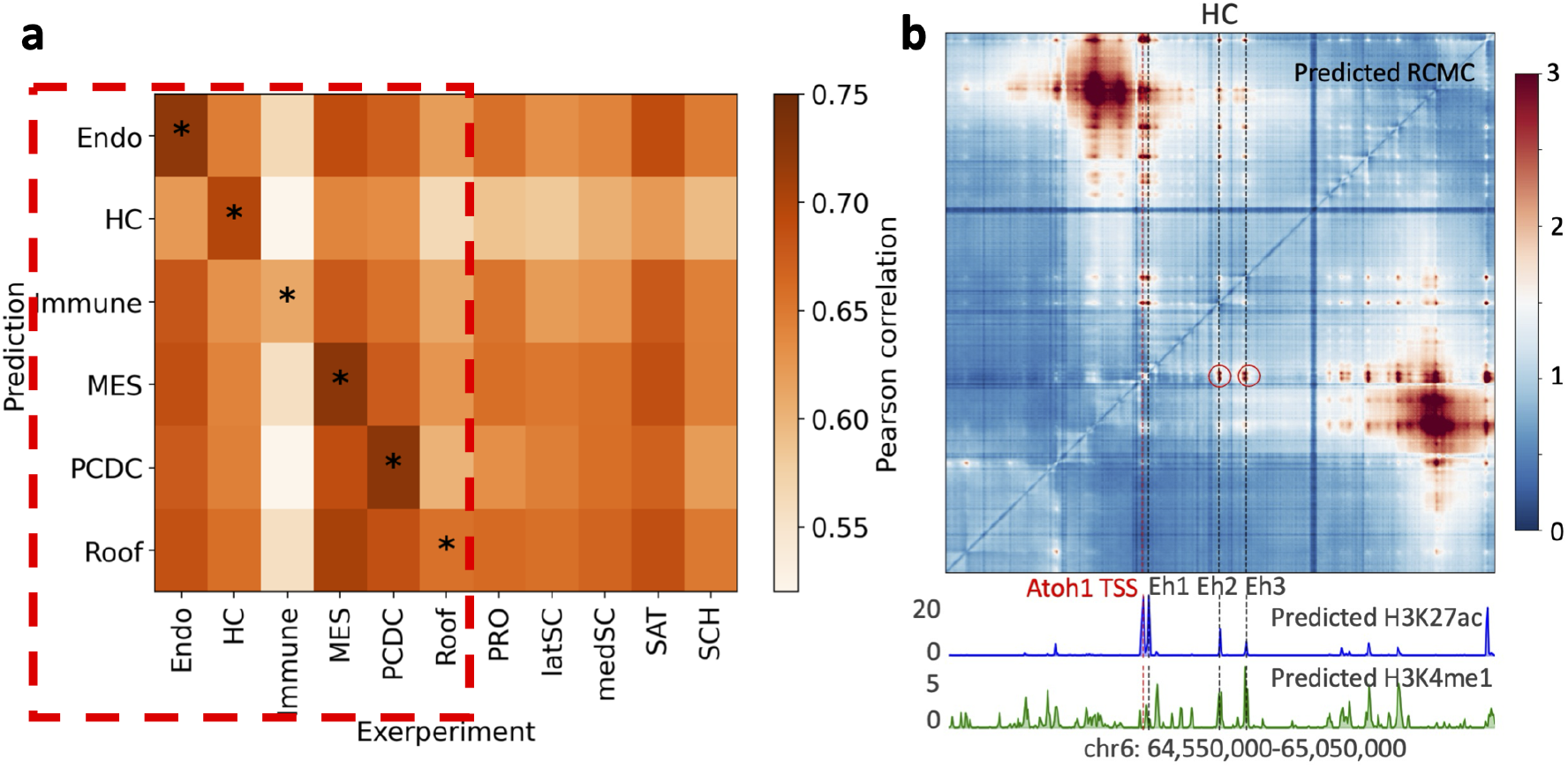
The model identified cell-type specific transcription regulation insights with a mouse inner ear dataset. **a**, The scATAC-seq data were aggregated into six pseudo-bulk profiles based on cell types (labeled on the y-axis) to predict RNA expression, while experimental scRNA-seq data were grouped into 11 cell types (labeled on the x-axis). The heatmap shows Pearson correlation coefficients between the experimental scRNA-seq and the predict RNA expression. For each experimental cell type (columns), the highest correlation is observed with the predicted expression of the corresponding cluster. **b**, The predicted RCMC contact map, H3K27ac, and H3K4me1 signals in HC cell types at Atoh1 region are displayed. Histone mark peaks are identified at three enhancer loci in HC, with loop patterns involving two distal enhancers highlighted by red circles.

To further evaluate the model’s utilities, we used postnatal day 2 mouse inner ear HC and Pillar Cell/Deiters’ Cell (PC/DC) ATAC-seq data [59] as input and predicted RCMC contact maps and histone marks (H3K4me1 and H3K27ac) at the Atoh1 locus. This locus is well investigated by a recent study that identified three enhancers associated with Atoh1 in hair cells [61]. The predicted histone marks, H3K4me1 and H3K27ac, exhibited strong signals at all three enhancer regions, and the predicted O/E normalized contact maps revealed clear contact patterns between the Atoh1 TSS and enhancers associated with Atoh1 in sensory HC [61] (Figure 5b). These predicted patterns align well with the identification of three active enhancers [61]. In contrast, the general model predicted a silencer at the thrid enhancer locus in PC/DC cell type characterized by low H3K27ac and H3K4me1 signals and the absence of a predicted chromatin loop with Atoh1 (Figure S18). This prediction is consistent with the study [61].

## Discussion

In this work, we developed a general AI model using a multi-task architecture to incorporate diverse genomic knowledge within a single model, by training on an extensive collection of genomic sequencing datasets [62, 23, 63, 40, 22, 64, 65, 66, 34, 67, 18, 68, 69]. The model is enriched with a wide array of epigenomic features and experimental assays, including mRNA, nascent RNA, ultra-high-resolution chromatin organization, and enhancer activity across diverse cells and tissues. Our model utilizes a supervised training approach, not only because of the availability of extensive sequencing data but also in response to observations that current DNA foundation models, which primarily use self-supervised pre-training techniques [7, 8, 9, 70], do not show superior performance compared to supervised methods trained from scratch [8, 71, 72], such as Basset [73], Enformer [3], SEI [4], and DeepSTARR [5], even though these foundation models still require task-specific fine-tuning. Moreover, the representation learning in these foundation models is often ineffective across diverse genomic tasks [49].

Our model accurately predicts various types of nascent RNA assays, offering a powerful tool for transcription analysis. By integrating ATAC-seq data, our model enables the prediction of nascent RNA profiles in single-cell pseudobulk and tissue data, where experimental measurements are often challenging to obtain. This capability allows our model to serve as a computational substitute for transcriptional analysis across both gene and enhancer regions. While our model provides valuable insights, further improvements in resolution are necessary, particularly for capturing enhancer activity. Many enhancers are short in length, making it difficult to precisely measure their transcriptional output. Enhancing the model’s resolution will improve its ability to distinguish fine-scale transcriptional features, ultimately leading to a more detailed understanding of enhancer-mediated gene regulation.

Predicting diverse genomic modalities in mouse species is crucial for understanding mouse-specific regulatory mechanisms. We adapted a general human model to develop the first general model for mouse species that can predict diverse genomic modalities from ATAC-seq data and generalize to unseen cells and tissues. The general model training techniques also demonstrate superior performance on region-capture Micro-C data, which is limited to a small number of genomic regions. However, due to potential distribution shifts caused by experimental differences in both the input ATAC-seq and the output genomic modalities between human and mouse, we leveraged transfer learning and fine-tuning strategies. While some sequence-based methods have shown advantages by concurrently training on human and mouse genome sequences [74, 75], we plan to explore models trained on both species in future. These efforts represent a step toward a more comprehensive understanding of species-specific genomic regulation, furthering comparative studies between human and mouse. Beyond mouse models, future efforts to develop general models for other species could provide valuable insights into evolutionary and functional genomics across diverse organisms.

## Methods

### Model architecture

The model features a multi-task architecture comprising three key components: a task-shared local encoder, a task-shared global encoder, and task-specific prediction heads (Figure 1a). The model inputs consist of a 600kb DNA sequence alongside corresponding ATAC-seq data. Similar to EPCOT, we segment the 600kb sequence into 1kb genomic bins, which is analogous to the patchification strategy used in computer vision [76]. Then, we pad each 1kb bin with 300bp sequences upstream and downstream as flanking regions. Each padded 1kb bin undergoes initial processing by the task-shared local encoder, which is designed to extract local sequence features. The local encoder comprises *three* blocks (Figure S4b). In the first two blocks, the DNA sequence and ATAC-seq data are processed separately to accommodate different receptive field requirements—narrower for motif patterns in DNA and broader for the chromatin landscape revealed by ATAC-seq (Figure S4b,c). For the DNA sequence *x*_*s*_, each block consists of two convolutional layers followed by a max-pooling layer, while for the ATAC-seq *x*_*a*_, the processing involves two convolutional layers, a bidirectional LSTM (Bi-LSTM) layer [77] to increase the receptive field, and a subsequent max-pooling layer. The outputs from both streams are then summed to provide a unified feature representation for subsequent layers, as shown in the following equations:

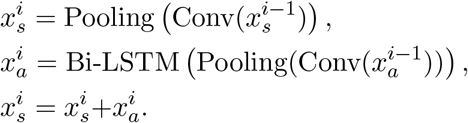

Then the third block which contains two convolutional layers and one transformer encoder layer to obtain the local features from the outputs of the first two blocks

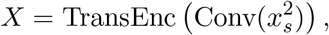

where *X*∈*R*^*n×d*^, and *n* indicates the number of extracted local features for each 1kb bin and *d* is the embedding size.

The features extracted by the local encoder are input to a multi-head global attention pooling layer to obtain a vector of local sequence representation *ϕ*∈*R*^*d′*^ for each local genomic bin. The pooling mechanism is defined as follows:

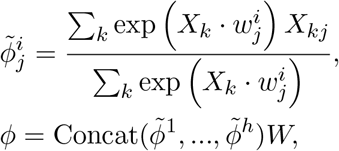

where *w*^*i*^∈*R*^*d×d*^ is the learnable weight matrix for *i*-th head, *W* ∈*R*^*hd×d′*^ is a weight matrix that integrates the outputs from different heads.

The local sequence representations of 600 1kb bins are then processed by a task-shared global encoder. The encoder comprises several convolutional layers followed by seven transformer encoder layers, updating the local sequence representations into global ones. The convolutional layers capture local interactions among different genomic regions, updating the representations *ψ*∈*R*^*N×d′*^, where *N* is the total number of genomic bins, set here as 600. Subsequently, six transformer encoder layers learn the long-range interactions among these regions

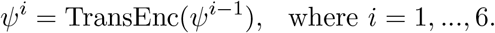

The attention logits within these layers are computed similarly to those in Enformer [3]

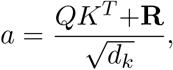

where **R** are relative positional encodings, and *Q* and *K* are query and key matrices used in self-attention [78], with *d*_*k*_ denoting the key dimension.

Next, the 1D global sequence representation *ψ* is transformed into 2D representations Ξ∈*R*^*N×N×d′*^ for chromatin contact map prediction (Figure S4e). These 2D representations are further processed by a contact map prediction (COP) head consisting of dilated convolutional layers:

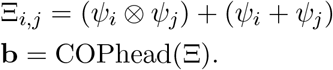

The 2D representations 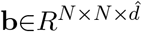 are leveraged for predicting 1kb-resolution chromation contact maps, including Micro-C and Intact Hi-C, through a linear layer. The representations also serve as a bias term in the calculation of attention logits in the last transformer encoder layer, which incorporate the chromatin interaction information to update the global representation

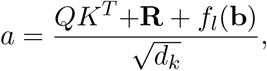

where *f*_*l*_ is a linear layer.

Finally, the last transformer encoder layer further refines the global sequence representations. Task-specific prediction heads, which are various linear layers, are then applied to the representations to predict specific genomic modalities.

### Mouse general model training

To effectively train the proposed general model to accommodate diverse genomic modalities, we implemented three strategies [79]. First, *task scheduling* is used to determine the orders in which tasks are introduced during the training. Second, *task weighting* is employed, where loss weights are assigned to the gradients of various tasks to balance them. Third, due to missing experimental data for some genomic modalities in specific tissues and cells, *partial label learning* is adopted.

For task scheduling, we implemented a curriculum learning approach [80], gradually introducing different genomic modalities into the training based on a pre-defined curriculum (Figure S1). Our approach involves two training schedules. In the first schedule, we include main courses that cover various genomic knowledge areas including epigenome, mRNA, nascent RNA, and high-resolution chromatin interactions. Training begins with these courses, progressing from tasks that rely primarily on local sequence information to those requiring long-range interaction information. We start with ChIP-seq data, which assesses protein binding on short DNA sequences, serving as a fundamental task and the pre-training step in our curriculum. Afterwards, we incorporate Bru-seq, BruUV-seq, and BruChase-seq [67, 81], which capture nascent RNA transcripts including enhancer RNAs and pre-mRNAs. Subsequently, we introduce CAGE-seq and RNA-seq, which measure mRNA and require information on distal enhancers due to their reliance on long-range interactions. Then, the curriculum progresses to include 2D chromatin interaction data, beginning with ChIA-PET [82], which provides protein-specific chromatin interaction information, and then advancing to high-resolution chromatin contact maps (e.g., Micro-C). After these main courses, additional genomic modalities are integrated into the training as extra courses. In the second training schedule, the model’s training initially focuses on cell line data, with tissue-specific data incorporated after all modalities in cell levels have been included.

Each component of the general model is gradually trained in the training schedule. Initially, we pretrain the local encoder using a method similar to EPCOT for binary activity classification on ChIP-seq data, where an encoder-decoder structure is employed during pre-training. After pre-training, the decoder is discarded, and only the local encoder is retained for subsequent stages. Then we include the global encoder except the last transformer encoder layer in the model training, and gradually train the specific prediction heads as different modalities incorporated in the training. After the 2D chromatin contact maps being predicted, the last transformer encoder layer which uses the chromatin interaction information in the self-attention layer is included and trained to update the predictions. The training schedule introduces a new course every 10 epochs, optimizing with the AdamW optimizer [83] at an initial learning rate of 5e-5, and employing MSE loss for all genomic modalities. Once all courses in both cells and tissues are integrated and trained, we shift to a reduced learning rate of 1e-5 for fine-tuning. The general model is initially trained in parallel on five NVIDIA A40 GPUs with 48GB memory, and then fine-tuned on five H100 GPUs.

For task weighting and partial label learning, we employ a *fixed weighting* approach and a *zeroing loss* strategy, respectively. This approach is simple yet effective in balancing training efficiency and model performance [79, 84, 85]. We maintain fixed loss weights across all tasks but increase the weights for each new course introduced during the training. Additionally, for genomic modalities that lack experimental data in certain cells or tissues, we zero the losses. Other approaches for partial label learning, such as developing teacher models on fully labeled data [11] for different modalities, exist. However, in our case, multiple modalities of data are missing in tissues, making it unsuitable to train a cell-level-only teacher model to generate pseudo labels for training in tissues. We thus chose a simple zeroing loss strategy.

### Training mouse general model

For training the mouse model, we adapt the pre-trained human general model by directly transferring and fine-tuning it, without employing the curriculum learning strategy. However, we continue to use fixed weighting and zero loss training approaches. Unlike other genomic modalities that are genome-wide, region-capture Micro-C (RCMC) data [18] is only available in five specific genomic regions totaling 5Mb. To effectively utilize the RCMC data, we upsample these regions by replicating them across all GPUs used in the training process, rather than distributing them separately to each GPU. As the mouse general model is also trained on five A40 GPUs, the samples associated with RCMC are effectively sampled five times. This approach allows the RCMC task to share encoders with other tasks, and only needs to train a specific prediction head for the RCMC task, thus preventing the model from overfitting on the limited data. The training parameters and techniques remain consistent with those used for the human general model, including the use of MSE loss and the AdamW optimizer. The training starts with an initial learning rate of 1e-4, which is subsequently reduced to 1e-5 to refine the model.

### Classification of eQTL in putative non-coding regions

To differentiate causal from non-causal variant-gene pairs, we employ a specialized architecture based on a pre-trained general model. We focus on fine-tuning only the local encoder, a small part of the entire model, to classify variants. This fine-tuning is predicated on the hypothesis that an eQTL may alter local sequence features and influence chromatin interactions with the target gene. The local encoder processes both reference and alternative sequences along with ATAC-seq within a 1.6kb region centered around the variant, using a Siamese structure (Figure 3a). We calculate differences in feature maps from each block of the encoder between the reference and alternative alleles, then aggregate these differences into a local feature difference vector 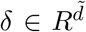. Then the entire general model remains frozen, generating a 2D representation **b**, where the element *i, j* corresponds to interactions between a variant *i* and a gene TSS *j*. Additionally, a prompt module integrates extra information such as variant scores from the Borzoi model [6] and gene types (0 for protein-coding, 1 for lncRNA, and 2 for others) (Figure S12a). This module also processes local features from the variant and gene TSS sequences, which are then combined with gene type embeddings through linear layers to update the 2D interaction feature **b**_*ij*_, and the variant score features are added to the local feature difference vector *δ*. Next, a gated function is then employed to aggregate these features, where

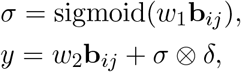

with 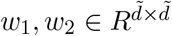 as trainable weight matrices, and ⊗ indicating element-wise multiplication. Finally, a linear classifier is applied to *y* to predict the probability of a variant-gene pair being causal. In the training phase, we freeze the parameters of the local encoder for the first five epochs. Subsequently, we fine-tune the encoder for this specific task.

To investigate the potential regulatory impact of non-coding variants on gene expression, we integrate eQTL datasets and ATAC-seq experiments. We download eQTL datasets from the eQTL catalog [13], which has uniformly processed eQTLs from various studies and we leverage variants quantified at the gene expression and exon expression levels in this dataset. For the identification of putative non-coding variants, we overlap these variants with ATAC-seq peaks. This involves downloading cell/tissue-specific ATAC-seq files and merging them when experiments include multiple donors. We then employ the MACS3 peak caller [86] to identify peaks from the ATAC-seq profiles using the command: `macs3 callpeak -g hs -q 0.01 -f BAMPE`. Variants found within these peaks in specific cells or tissues are considered putative non-coding variants.

For the training data, we utilize these identified putative non-coding variants from 25 tissues and cells from the GTEx (v8) project [40], with available ATAC-seq or scATAC-seq data. For testing the model, we use variants from 13 different cell types across five additional studies (schmiedel 2018 [44], Schwartzen- truber 2018 [45], Alasoo 2018 [46], HipSci [47], and iPSCORE [48]), where corresponding ATAC-seq or scATAC-seq data is available. Similar to prior studies [3, 87], our training eQTL dataset categorizes variant-gene pairs as causal if they have a PIP score greater than 0.8, as determined by the SuSIE fine-mapping method [88], and as non-causal if their PIP score is below 0.1. For the testing eQTLs, we adopt a more stringent selection criterion, identifying positive variants with PIP scores above 0.9 and negative variants with PIP scores below 0.05. In both the training and testing set, the negative samples are randomly selected to maintain a positive to negative ratio of 1:5. Similar to the approach used in Enformer, for each positive variant-gene pair, we first select negative pairs corresponding to the same gene.

We apply five-fold cross-validation to the GTEx training dataset. In each fold, we exclude any variants from the test set that appear in the training set, even if they are associated with different tissues or cells, to ensure complete independence in testing. Through this process, all variants in the test set are tested. We selected two state-of-the-art sequence-based models as baselines: Enformer [3] and Borzoi [6]. Both models utilize random forest classifiers to discern causal eQTLs from non-causal ones based on variant scores—Borzoi employs ‘L2’ scores, while Enformer uses ‘sum’ scores. Additionally, we have introduced a crucial feature, the distance from the gene TSS to the variant scores [87]. Consequently, Enformer’s variant score vectors comprise 5314 features, and Borzoi’s contain 7612 features.

### LentiMPRA prediction

For lentiMPRA prediction, we utilized data from three cell lines in the study [14]. We first converted the data from the hg19 to the hg38 genome assembly. Only elements that have regulatory scores on both forward and reverse strands are retained. Following the study [49], we used the forward sequence to predict regulatory scores on the forward strand and the reverse complement sequence to predict scores on the reverse strand, but we ensured that the forward and reverse sequences of the same element were placed in the same training, validation, or testing set to prevent information leakage. As the sequences involved were short, the prediction was achieved by utilizing and fine-tuning only the local encoder, which takes an input sequence of 1.6 kb. Each element is centered within the input, with upstream and downstream sequences added as flanking regions. Additionally, convolutional and linear layers were applied on top of the embeddings output from the encoder (Figure S11).

In cross-cell type lentiMPRA prediction, we randomly split elements from K562 and HepG2 into training and validation sets using an 80:20 ratio. The model was subsequently tested on all elements from WTC11. In within-cell type prediction, we trained the model separately for each cell line, adhering to a training, validation, and testing distribution of 70%, 10%, and 20%, respectively.

Three baseline models, MPRAnn, MPRALegNet [14] and ResidualBind [49, 89] were employed, which are end-to-end deep learning models capable of integrating cell-type-specific ATAC-seq data through minor architectural modifications (the kernel size of the first convolutional layer is increased from four to five). To integrate cell-type-specific ATAC-seq with DNA sequence, the barcode sequence was replaced with a 15 bp reference sequence upstream and downstream as flanking regions, resulting in a 230bp DNA sequence. When ATAC-seq was incorporated, the rest of the architecture remained unchanged. Additionally, although ResidualBind utilized an exponential activation function after the first convolutional layer, we found it did not perform well with ATAC-seq input. Therefore, we replaced exponential activation with a ReLU activation function in our model. Both methods have been re-implemented in PyTorch.

### Enhancer-gene regulatory interaction prediction using perturbation data

We utilized the K562 CRISPR perturbation data collected in the study [15] for validation and retrained the general model without using the K562 cell line (see training cells and tissues in Figure S13a). The original dataset identified 472 enhancer-gene regulatory interactions in K562 cells. We used 451 of these for our analysis, covering a total of 6,461 element-gene pairs within our predictive range. To test whether the cell-type specific knowledge from our general model could improve ENCODE rE2G’s performance, we included either learned representations or predicted modalities as additional features, alongside the original ENCODE rE2G’s features. When using the general representations, we incorporated both the 1D sequence representations *ψ*, output from the final transformer Encode layer, of the gene TSSs and the element bins, and the 2D sequence representations **b** of the chromatin interaction between the gene TSS and the element bins. A random forest classifier, implemented using Scikit-learn’s default settings [90], was then applied to the *z*-score normalized representations. For efficiency, we opted not to use the default logistic regression employed by ENCODE rE2G. Additionally, when using the predicted modalities, we utilized the predicted signal values of the modalities at both the gene TSS and element bins, and integrated them into ENCODE rE2G’s features. A logistic regression classifier was then used to predict the regulatory interactions.

### TT-seq experiments and analysis

The TT-seq procedure largely followed our published method [22, 91]. Briefly, 5–8 million MCF10A cells were cultured in 10 cm dishes and treated with TGF-*β* (10 ng/mL, Pepertech 100-21-100UG) or a vehicle control for 24 hours, reaching approximately 70% confluency at the time of harvest. The cells were then incubated with 4-thiouridine (4SU) (Sigma T4509-250MG) at a final concentration of 700 *µ*M for 15 minutes at 37°C. Total RNA was extracted using TRIzol, and fragmented RNA was biotinylated with MTSEA-biotin-XX (Biotium, 90066-1) for 2 hours. Biotinylated 4SU-labeled nascent RNA was isolated using Streptavidin C1 beads (Thermo fisher 65002). RNA libraries were prepared with the VAHTS Universal V8 RNA-seq Library Prep Kit for Illumina (Vazyme NR605-02). Sequencing of the library was done by the Illumina NextSeq 550 system with 40bp PE mode (by UTHealth Cancer Genomics Core). TT-seq reads were trimmed with Trim Galore v0.6.10, aligned to the human genome (hg38) using STAR v2.7.3a using default parameters, and duplicate reads were removed with Picard. Reads were quantified by HOMER, and differential gene expression analysis was conducted by EdgeR using FDR 0.05 as cutoff of significance.

## Acknowledgment

This work was supported by the National Institutes of Health under grants R35HG011279, R03OD038390, R01DK129469, U01DK135017, U24DK138515, and U01HG011952.

## Data and code availability

The code and accession numbers for the genomic modalities used in this study are available in our GitHub repository https://github.com/liu-bioinfo-lab/general_AI_model. A tutorial for using our model is publicly available at https://epcotv2-tutorial.readthedocs.io/en/latest/. Additionally, our model’s web portal can be accessed at https://huggingface.co/spaces/luosanj/EPCOTv2, where users can easily predict diverse modalities.

## Supplementary information

### Data processing

All sequencing data used in our model is in the hg38 version for humans and the mm10 version for mice. Since our model utilizes sequencing data from diverse cells and tissues in the training, we aimed to remove distribution shifts during training. For the TF and histone mark ChIP-seq data, we utilized their signal p-values and performed *z*-score normalization. We then clipped the normalized signal values within the range (−2, 36) to exclude extreme values. For all expression data, including CAGE-seq, RNA-seq, Bru-seq, BruUV-seq, BruChase-seq, TT-seq, GRO-seq, GRO-cap, PRO-seq, and Net-CAGE, we followed the normalization method used in GraphReg [92] for cross-cell type prediction. We applied RPGC normalization [93] to generate the bigWig files with the binsize set to the resolution to be predicted: `bamCoverage --normalizeUsing RPGC --effectiveGenomeSize 2913022398 --binSize 1000`. Subsequently, we divided each signal value by the 95th percentile of non-zero values and performed an arcsinh transformation to preserve the original distribution. This approach avoids the distributional distortions that direct logarithmic or arcsinh transformations might introduce.

For Hi-C, ChIA-PET, and Micro-C contact maps, we used Juicebox [94] to derive the KR or O/E normalized contact maps. For KR normalization, we used Juicebox straw’s ‘SCALE’ normalization. The intact Hi-C and Micro-C contact maps at a 1kb resolution were log2 transformed and clipped within the range (−2, 10). Subsequently, the intact Hi-C data was processed using a Gaussian smoothing filter with a length of 5 and a sigma of 1, while the Micro-C data was smoothed with the same filter length but a sigma of 0.8. For the ChIA-PET data at a 5kb resolution, we applied a log2 transformation with a pseudocount of 1, followed by Gaussian smoothing with a filter length of 5 and a sigma of 1.

The ChIP-seq, CAGE-seq, RNA-seq, Bru-seq, BruUV-seq, BruChase-seq, GRO-cap, STARR-seq, and Intact Hi-C data were downloaded from ENCODE [23], while ChIA-PET and Micro-C data were sourced from 4DN. For data originating from different studies, GRO-seq was aligned using Bowtie2 [95], and both TT-seq and NET-CAGE were aligned using STAR [96].

**Fig. S1.**
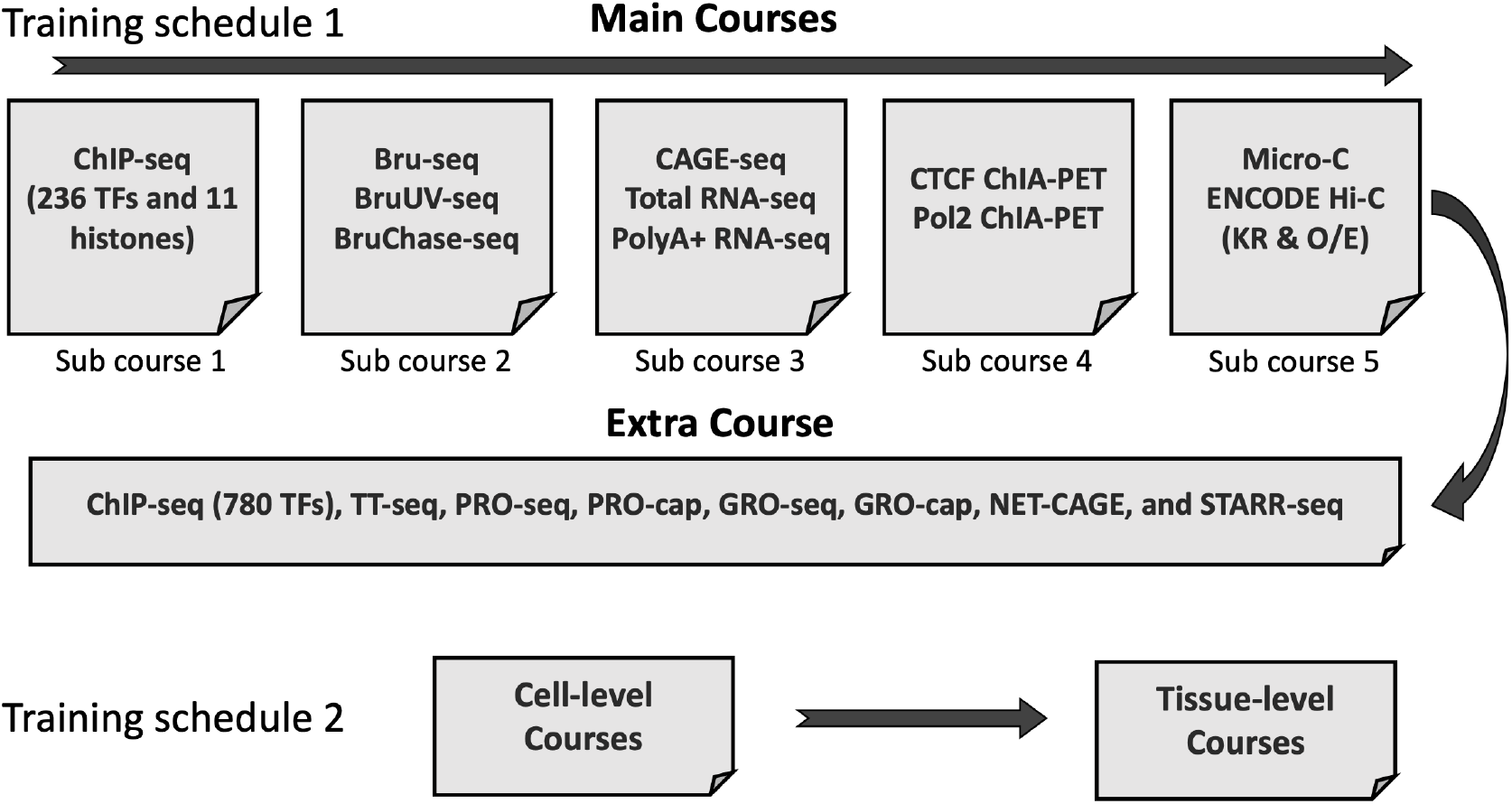
Two training schedules included in our general model’s curriculum learning framework. One schedule introduce the model from main courses to extra courses, and the other one introduce the model from cell data to tissue data.

**Fig. S2.**
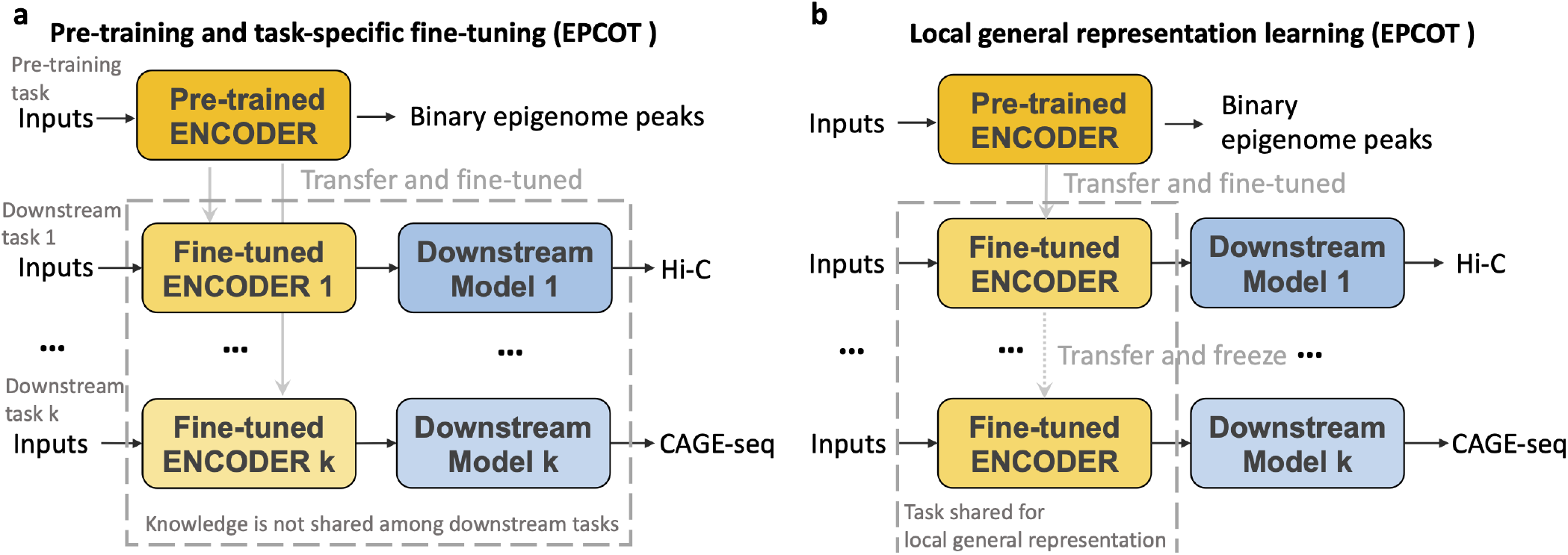
Strategies used in EPCOT model. **a**, EPCOT uses a pre-training and fine-tuning framework which is pre-trainined to predict binary peaks of epigenomic features, and then the encoder is separately transferred and fine-tuned in several downstream tasks and task-specific downstream models are trained. **b**, A simple strategy used in EPCOT for general representation learning. The pre-trained encoder in epigenomic feature task is further fine-tuned in Hi-C prediction task. Then this encoder is frozen to generate local general representations for other predictive tasks.

**Fig. S3.**
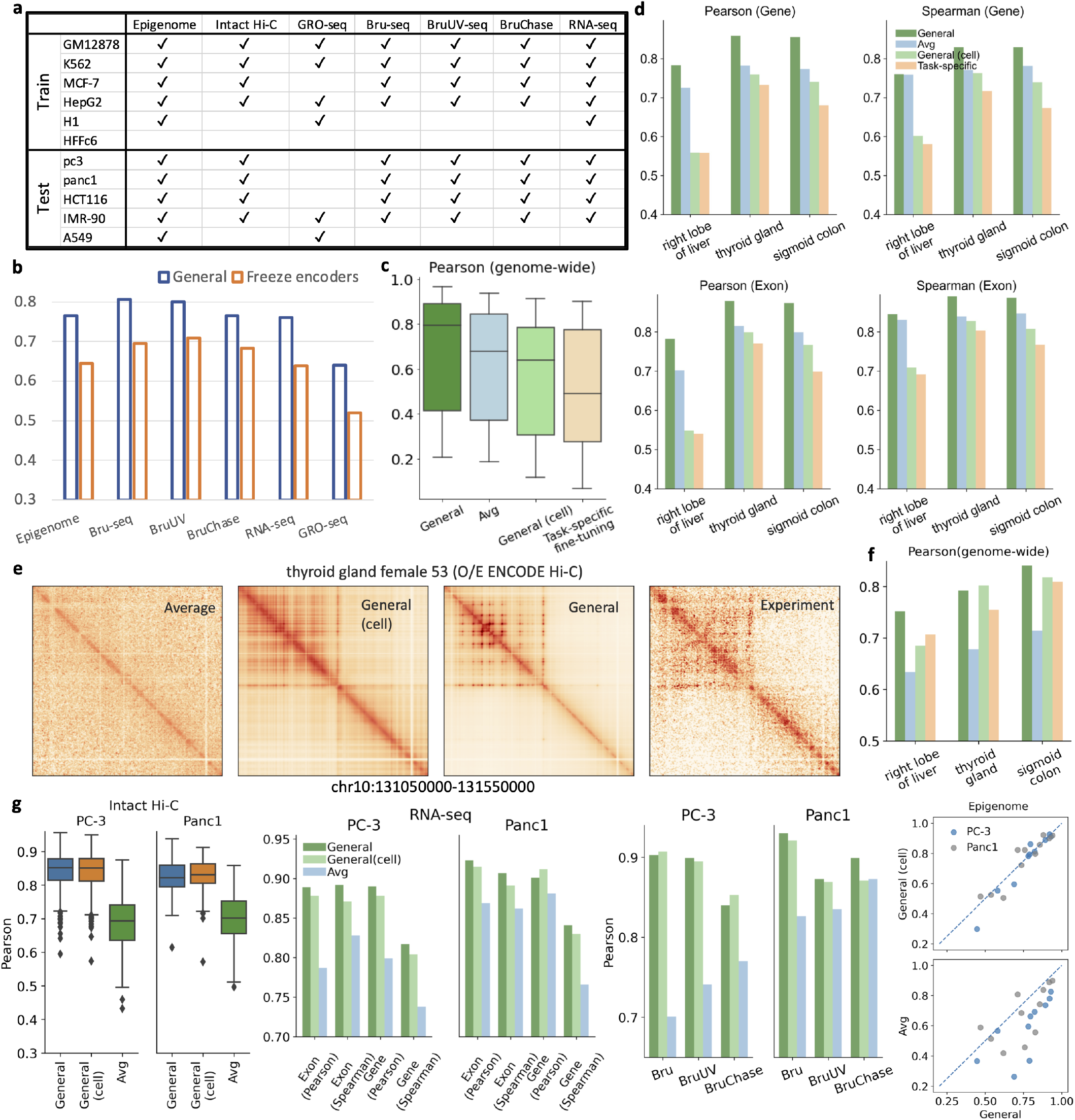
See captions on the next page. Evaluation of general model’s performance across diverse tasks. **a**, Availability of genomic modalities in training and testing cell lines used for evaluation. **b**, The general model outperforms the models by freezing both local and global encoders, pre-trained on epigenome prediction and then fine-tuned in Intact Hi-C prediction, which indicates better general representation learning. **c**, The general model incorporating multiple tissue data outperforms all other approaches including average signal experiment, the general model trained only on cell lines, and task-specific fine-tuning models, in predicting epigenomic features in three unseen tissues. **d**,**f**, The general model incorporating multiple tissue data outperforms all other approaches in the RNA-seq prediction task across genomic bins associated with the whole genome, genes, and exons in chromosomes 10 and 21. **e**, A region is illustrated to demonstrate that the general model predicts the Intact Hi-C contact map more accurately than the average Hi-C map. **g**, The general models, whether or not including multiple tissue data in the training, perform similarly across different tasks and both outperform average signal experiments.

**Fig. S4.**
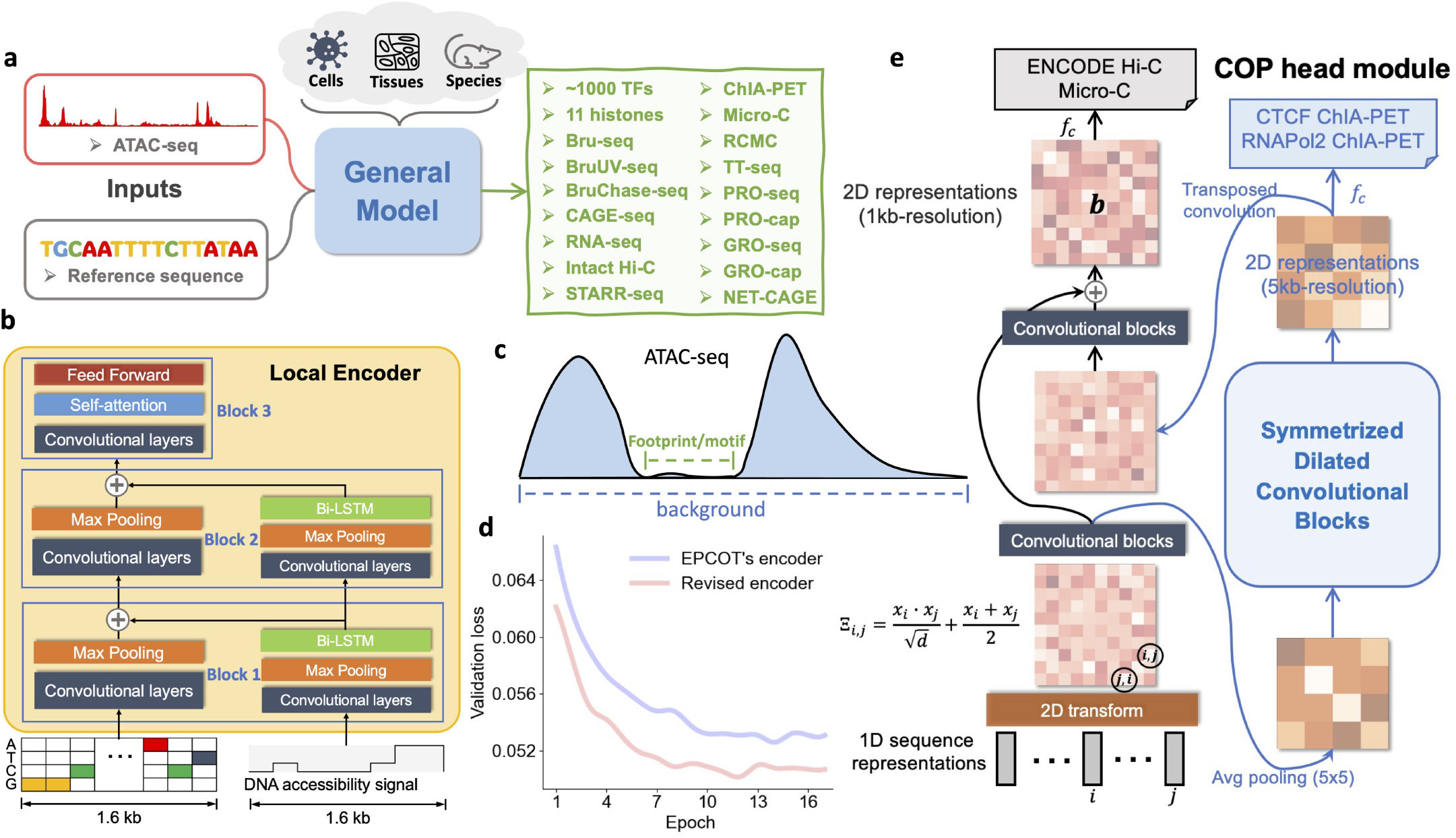
Model architecture illustration of the general model. a, An overview of the proposed general model. b, The model architecture of local encoder. c, An illustration of ATAC-seq experiments. d, The revised local encoder used in the general model achieves lower validation loss comparing to the encoder used in EPCOT. e, The architecture of the COP head module is depicted. 1D sequence representations are initially converted into 2D features Ξ, each representing chromatin contacts at 1kb-resolution. These features undergo convolution and pooling, followed by dilated convolutional layers to achieve 2D representations at a 5kb resolution, which are then utilized to predict ChIA-PET interactions. The representations are subsequently upsampled to 1kb-resolution 2D features, which are further refined through additional convolutional layers. A residual connection integrates these features with the pre-pooling 2D features, ultimately yielding refined 2D representations at 1kb resolution used to predict Intact Hi-C and Micro-C data.

**Fig. S5.**
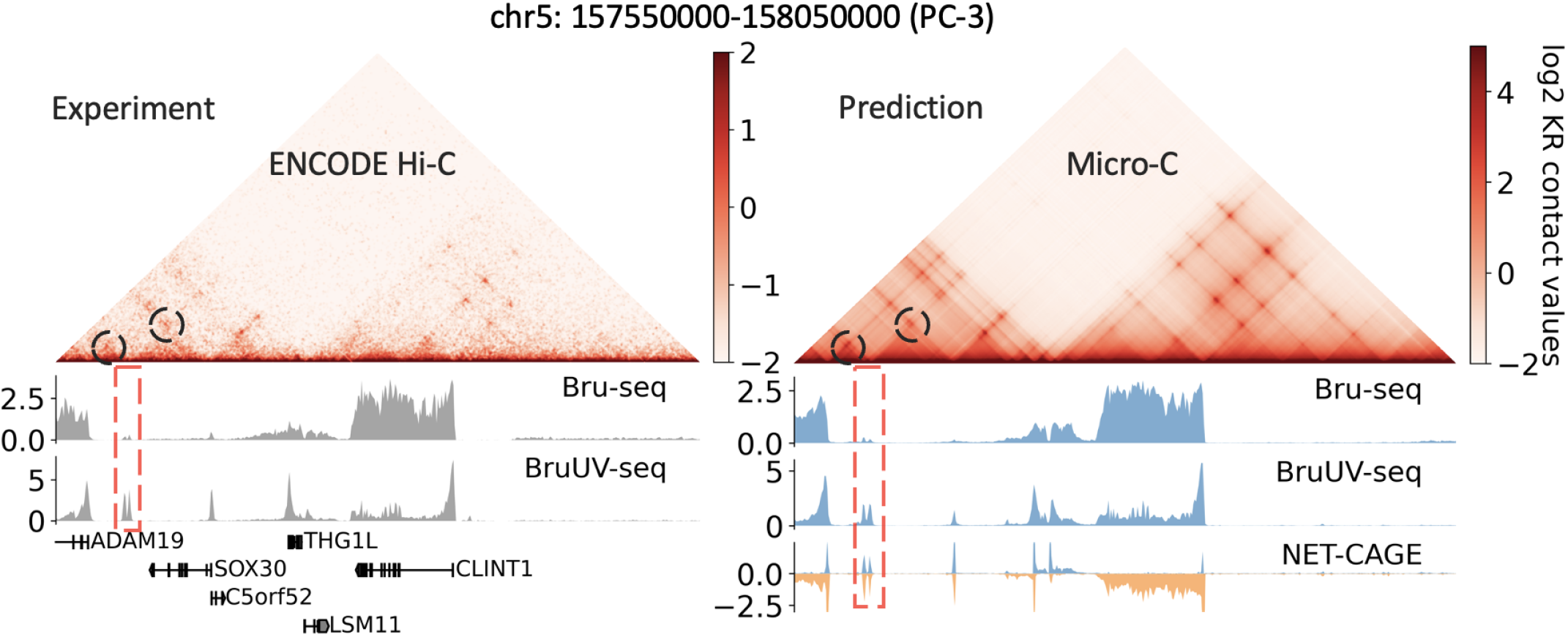
A 500 kb region in the PC-3 testing cell line demonstrates the accurate prediction of putative eRNAs (highlighted in red dashed boxes) and chromatin interactions with neighboring genes ADAM19 and SOX30 (highlighted in black dashed circles). These are validated by Hi-C, Bru-seq, and BruUV-seq experiments.

**Fig. S6.**
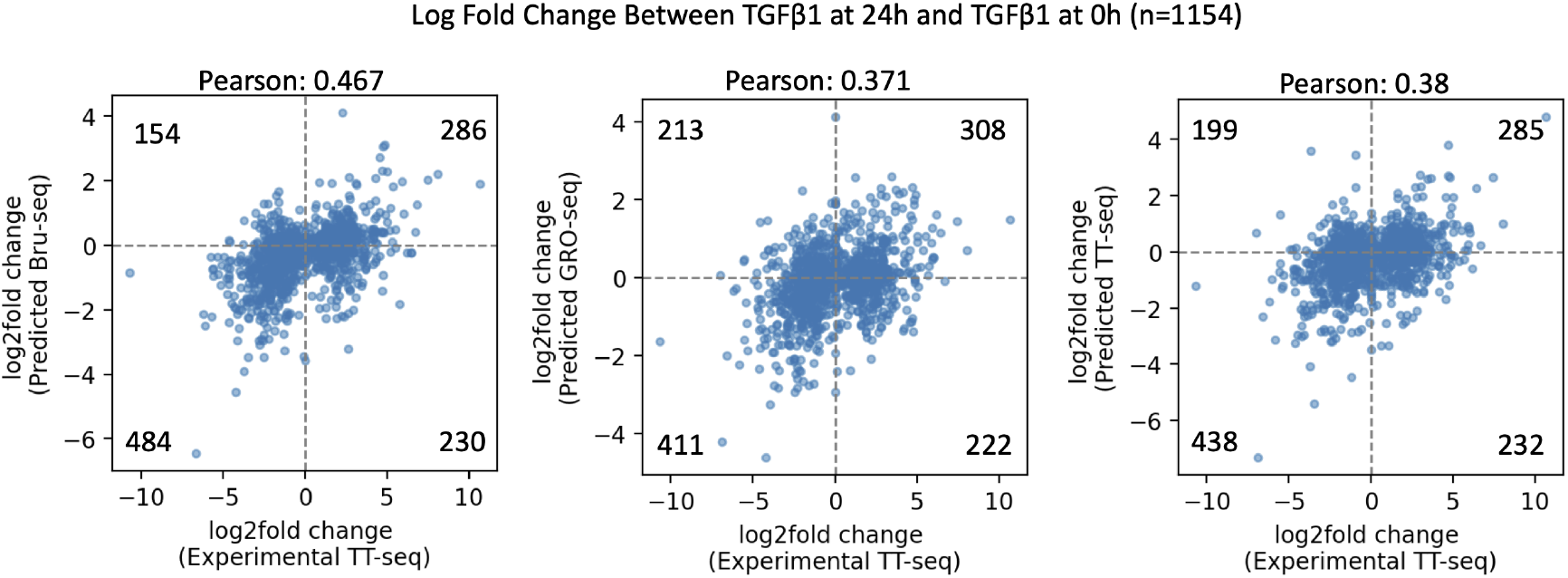
Correlation of predicted and experimental log2 fold changes in nascent RNA expression between TGF*β*-1 treatment at 24 hours and 0 hours across differentially expressed genes. Log fold changes predicted from Bru-seq, Gro-seq, and TT-seq are compared to the log fold changes obtained from experimental TT-seq. The results demonstrate the model’s ability to capture treatment-induced changes in gene expression. Numbers in each quadrant represent the count of data points in that region, corresponding to agreement or disagreement between predicted and experimental log2 fold changes. Having more Bru-seq data included in the training likely contributed to achieving the highest correlation score for Bru-seq.

**Fig. S7.**
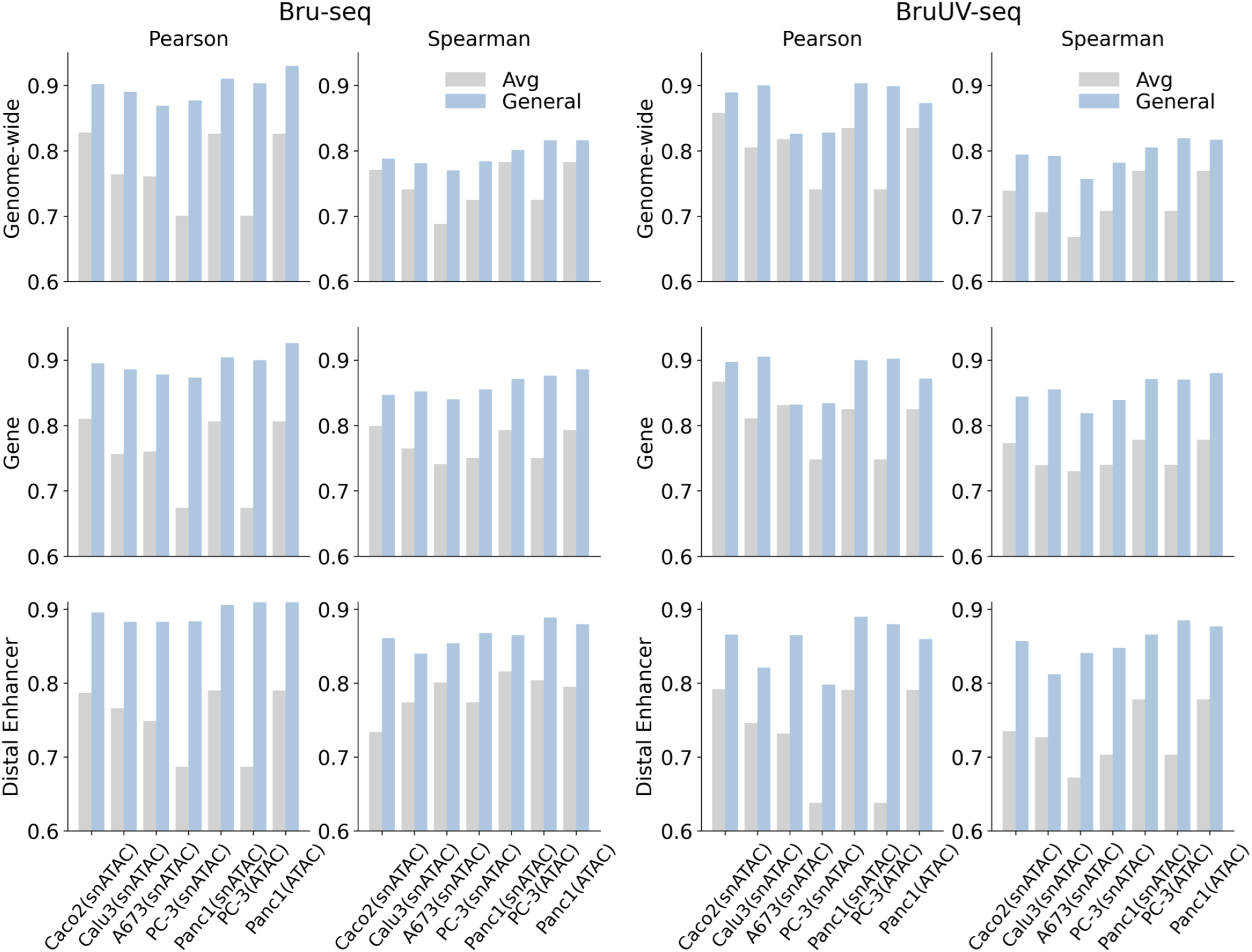
Prediction performance comparisons between the average signal of all training cell lines and the general model using snATAC-seq or ATAC-seq on Bru-seq and BruUV-seq. The general model outperforms the average signal experiment across genomic bins associated with the whole genome, genes, and candidate distal enhancer elements in chromosomes 10 and 21. For PC-3 and Panc1 cell lines, we use both snATAC-seq and ATAC-seq, while for the other three cell lines, we use snATAC-seq.

**Fig. S8.**
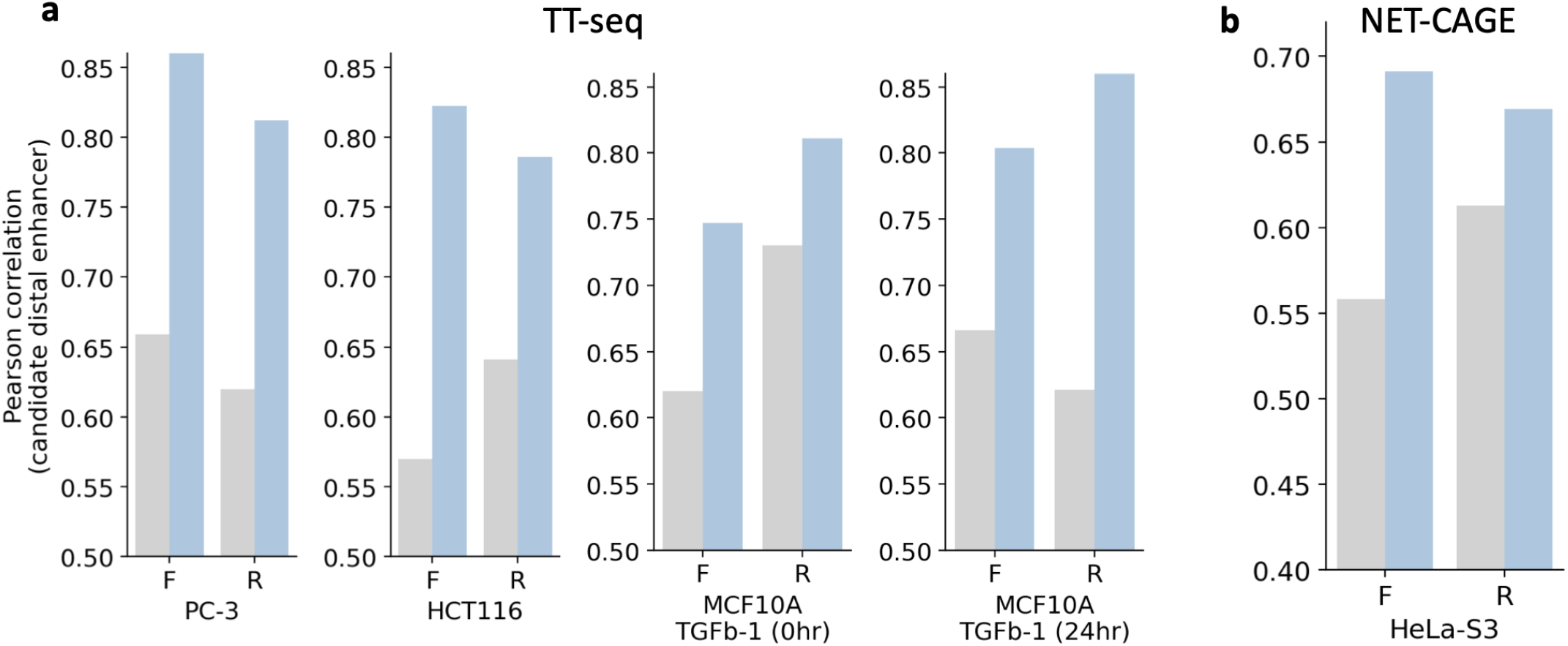
Prediction performance on TT-seq (a) and NET-CAGE (b) across candidate enhancer elements. Our model makes predictions more accurately than average experimental signals.

**Fig. S9.**
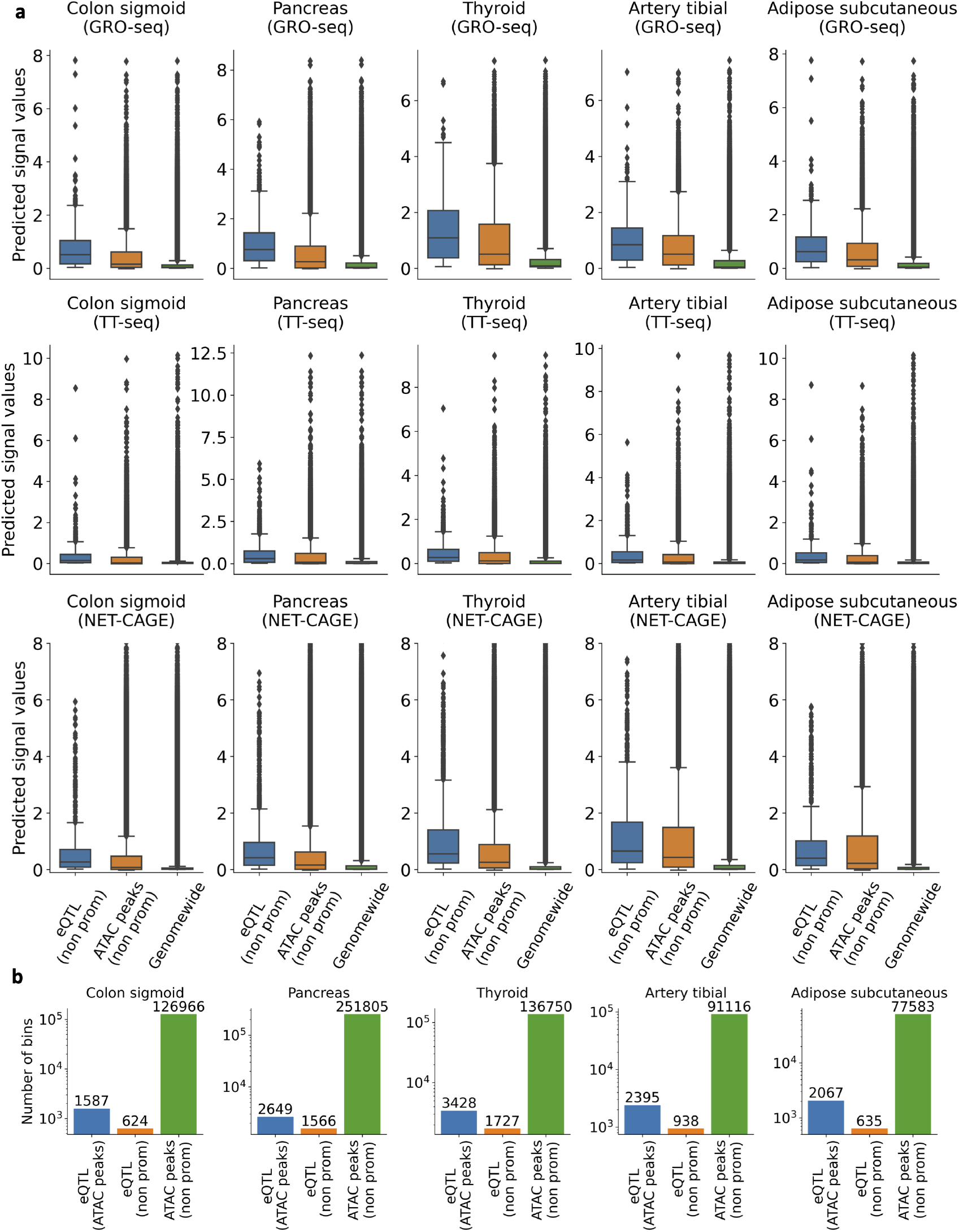
See captions on the next page. Predictions of GRO-seq, TT-seq, and NET-CAGE on five tissues. **a**, The predicted signal values for three types of regions are displayed. The first type associates with genomic bins containing eQTLs that have a PIP score greater than 0.1 and overlap with corresponding non-promoter ATAC-seq peak regions. For comparative analysis, we also use genomic bins that overlap with non-promoter ATAC-seq peak regions and all genomic bins as a background comparison. Generally, the model predicts higher signal intensities at eQTL loci. **b**, The numbers of bins associated with eQTLs in ATAC-seq peaks, eQTLs in non-promoter ATAC-seq peaks, and non-promoter ATAC-seq peaks in five tissues.

**Fig. S10.**
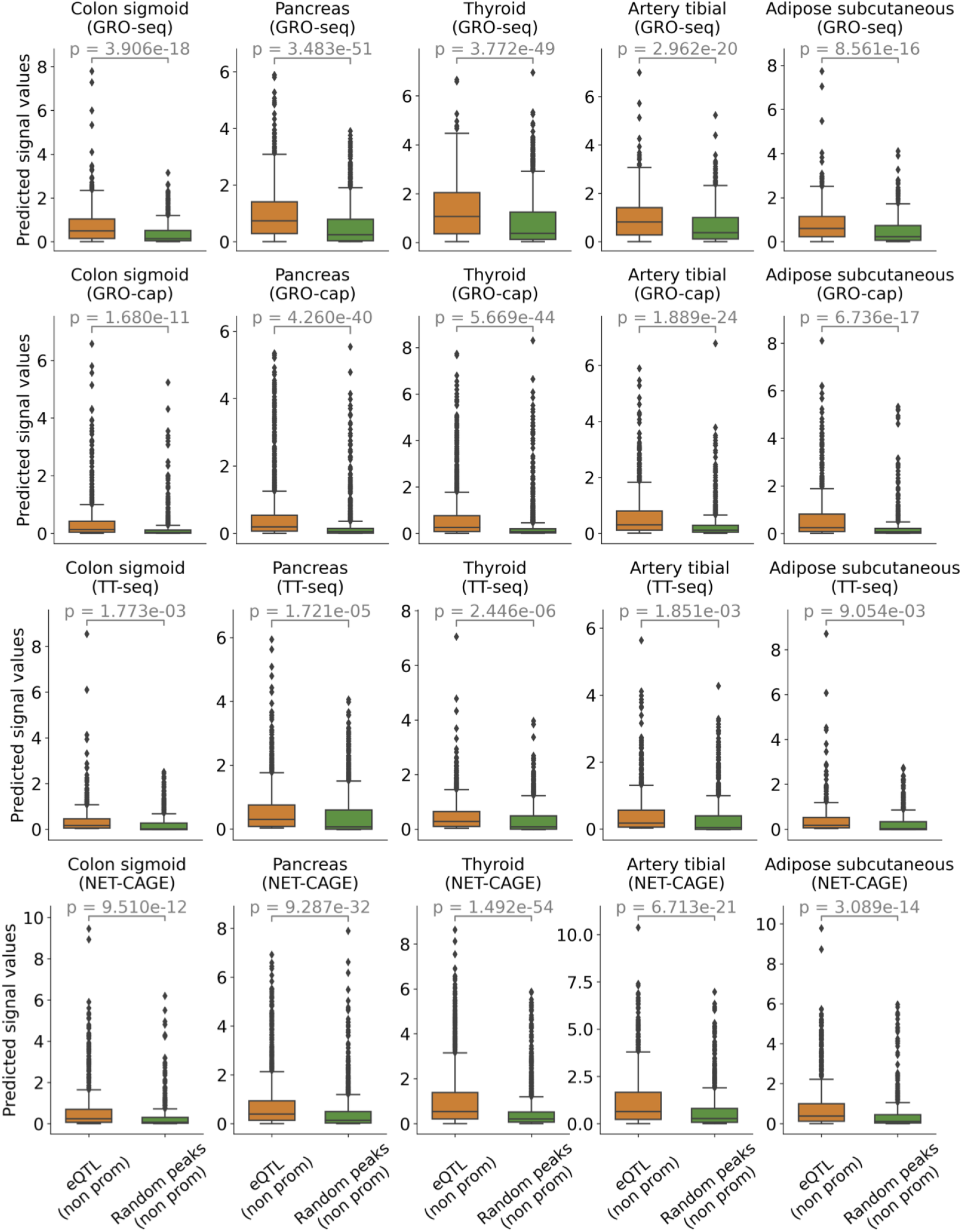
The general model predicts higher signal values in eQTL regions that overlap with non-promoter open regions compared to those in non-promoter open regions only. We first obtain the predicted GRO-seq, GRO-cap, TT-seq, and NET-CAGE signal values for genomic bins associated with eQTLs (PIP *>* 0.1) located in non-promoter open regions. We then randomly sample an equivalent number of genomic bins exclusively from non-promoter open regions and record their signal values. Subsequently, we perform a *t*-test on these predicted values to determine whether the signal values in eQTL regions are significantly higher.

**Fig. S11.**
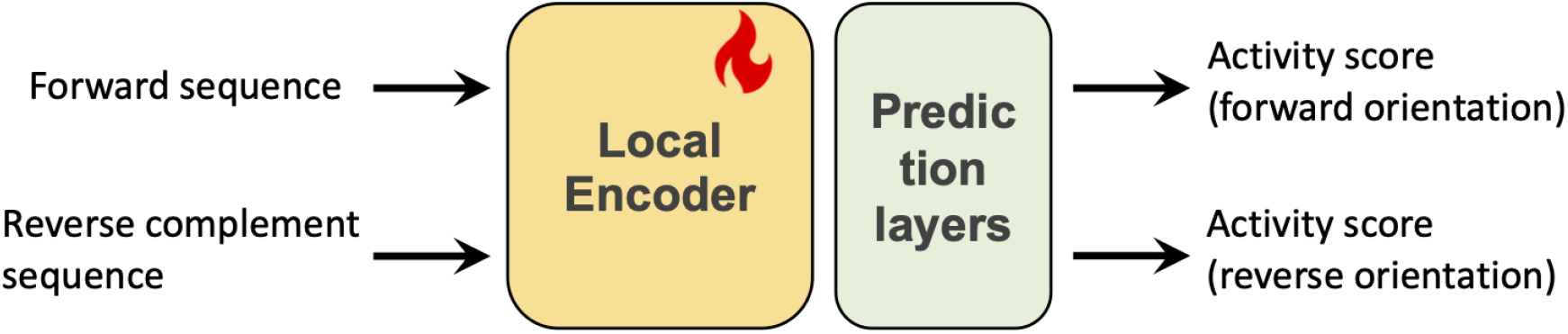
Description of the LentiMPRA prediction task. We utilized both forward and reverse complement sequences to predict regulatory activity scores in corresponding orientations. This prediction is achieved by fine-tuning the local encoder and training additional convolutional and linear layers.

**Fig. S12.**
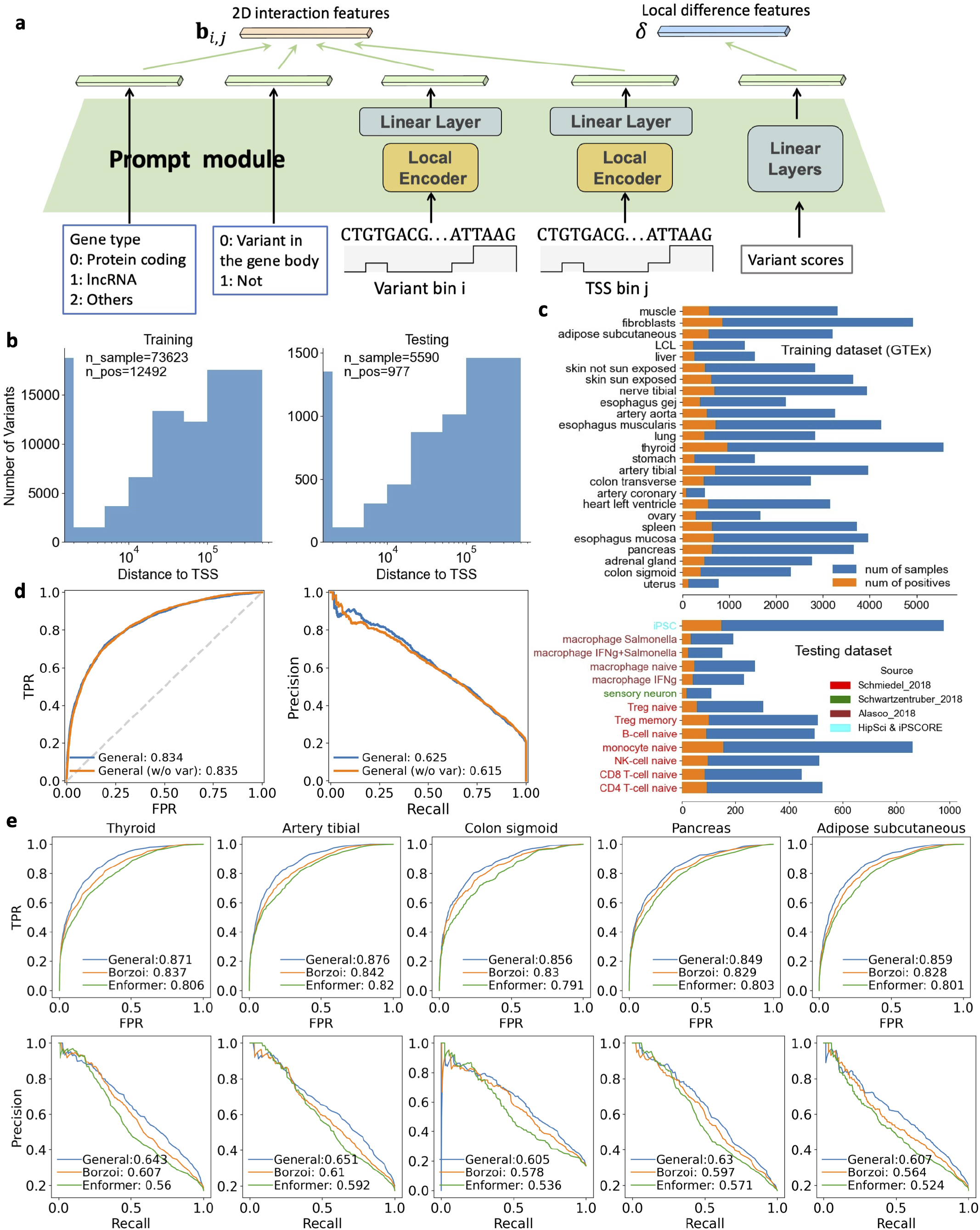
See captions on the next page. Using the general model to improve eQTL classification. **a**, The architecture of the prompt module introduces extra information. First, gene type information, including protein-coding, lncRNA, and other gene types, along with the position of the variant (whether in the gene body), are converted into learnable embedding vectors. Then, these vectors and two instance-dependent vectors generated from the variant bin *i* and TSS bin *j* using the local encoder from the general model, are integrated into the 2D interaction feature *b*_*i,j*_. Additionally, a Borzoi variant score vector is processed through linear layers and then incorporated into the local feature difference vector *δ*. **b**, Distribution of variants by distance to TSS in training and testing sets. The histograms show the number of variants as a function of their distance to the TSS. The number of samples, including those positive samples (causal eQTLs), is provided. **c** The number of samples, including the positive ones, in each cell or tissue from the training and testing datasets is provided. **d**, Our model achieves similar performance whether using Borzoi’s variant scores as extra information in cross-cell type and study eQTL prediction. **e**, Integrating the representations of the general model with Borzoi variant scores outperforms both Borzoi and Enformer models in five tissues. The AUROC and PR curves are visualized.

**Fig. S13.**
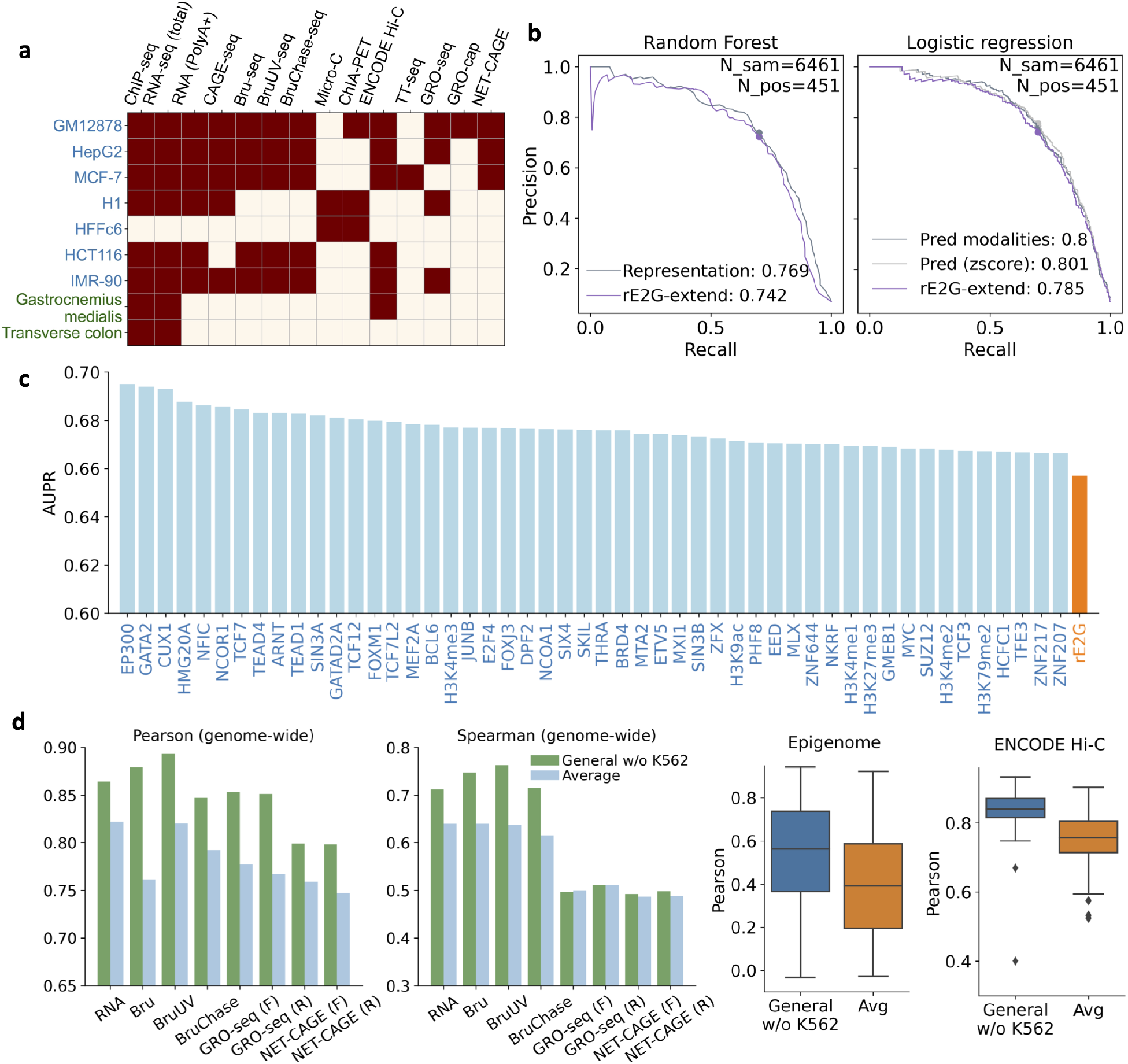
Evaluation results on the K562 CRISPR perturbation dataset. **a**, The cells, tissues, and genomic modalities used to retrain the general model were selected to exclude the K562 cell line, thus preventing information leakage during testing on the perturbation dataset. **b**, AUPR scores resulting from the introduction of individual predicted TFs to the ENCODE rE2G features are presented. The top 20 TFs showing the greatest improvement are displayed, with the original ENCODE rE2G score provided for comparison. **c**, Both the representations and predicted modalities improve the ENCODE rE2G-extend model. A random forest algorithm is applied to the *z*-score normalized representations and features used in the ENCODE rE2G-extend model. Logistic regression is employed for the predicted modalities, both with and without *z*-score normalization, along with the ENCODE rE2G-extend features. The predicted modalities include RNA-seq, CAGE-seq, GRO-seq, GRO-cap, NET-CAGE, and O/E and KR normalized intact Hi-C and Micro-C contact maps. Note that predicted histone marks and EP300 are not included, as their experimental data has already been incorporated into the ENCODE rE2G-extend features. d, Prediction performance comparisons are shown between average signals using data from Figure 1e, excluding K562 data, and the general models using only K562 ATAC-seq. The general model consistently achieves better performance.

**Fig. S14.**
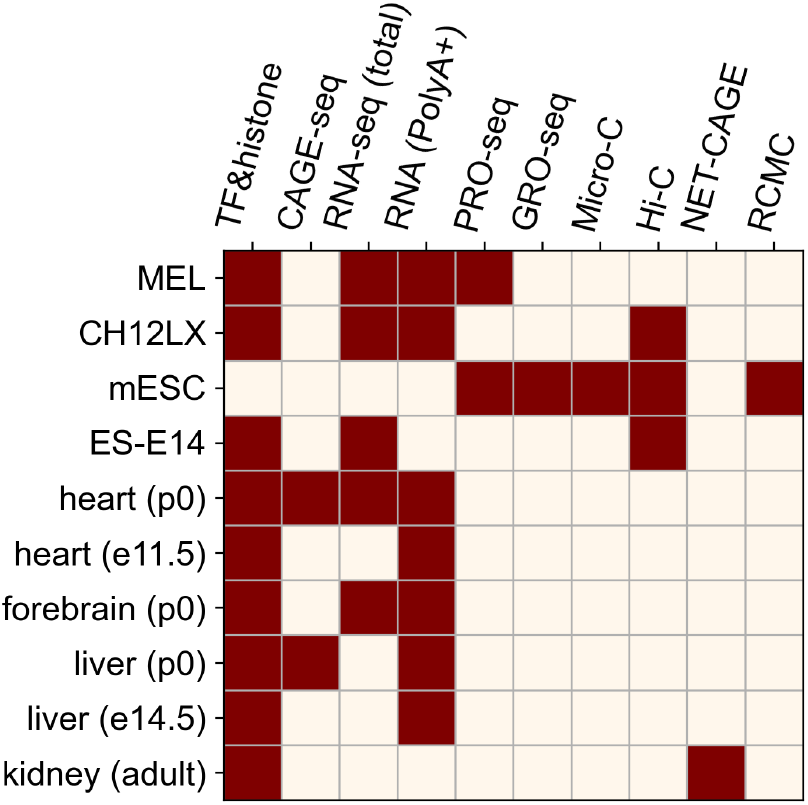
Overview of the cells and tissues as well as the modalities used in the training of mouse general model.

**Fig. S15.**
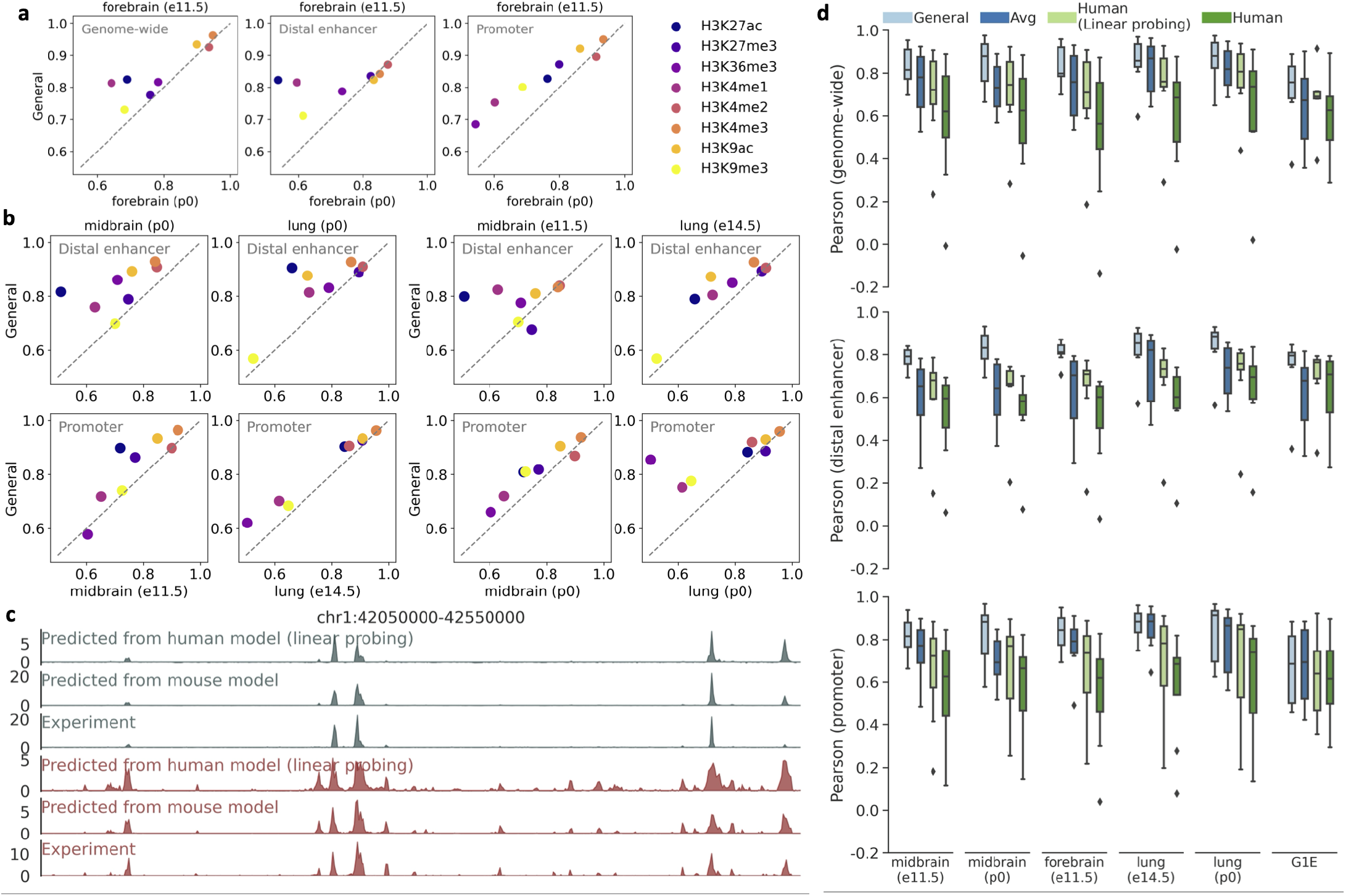
Prediction performance on mouse histone mark prediction. **a**, The general model predicts histone marks in forebrain embryos more accurately than using the experimental histone marks from forebrain postnatal across the genome, distal enhancers, and promoter regions in chromosome 1. The Pearson correlation scores are reported. **b**, The general model more accurately predicts histone marks in the midbrain and lung at one developmental stage compared to using their experimental embryonic histones from another developmental stage, across distal enhancer and promoter regions. **c**, A region to show that mouse model predicts histone marks more accurately than the human model using the linear probing method. **d**, Comparison of prediction performance among the mouse general model, average signals of all training cells and tissues, and the human model with or without fine-tuning of the last linear layer. The mouse general model consistently achieves better performance across the genome, distal enhancer, and promoter regions in chromosome 1.

**Fig. S16.**
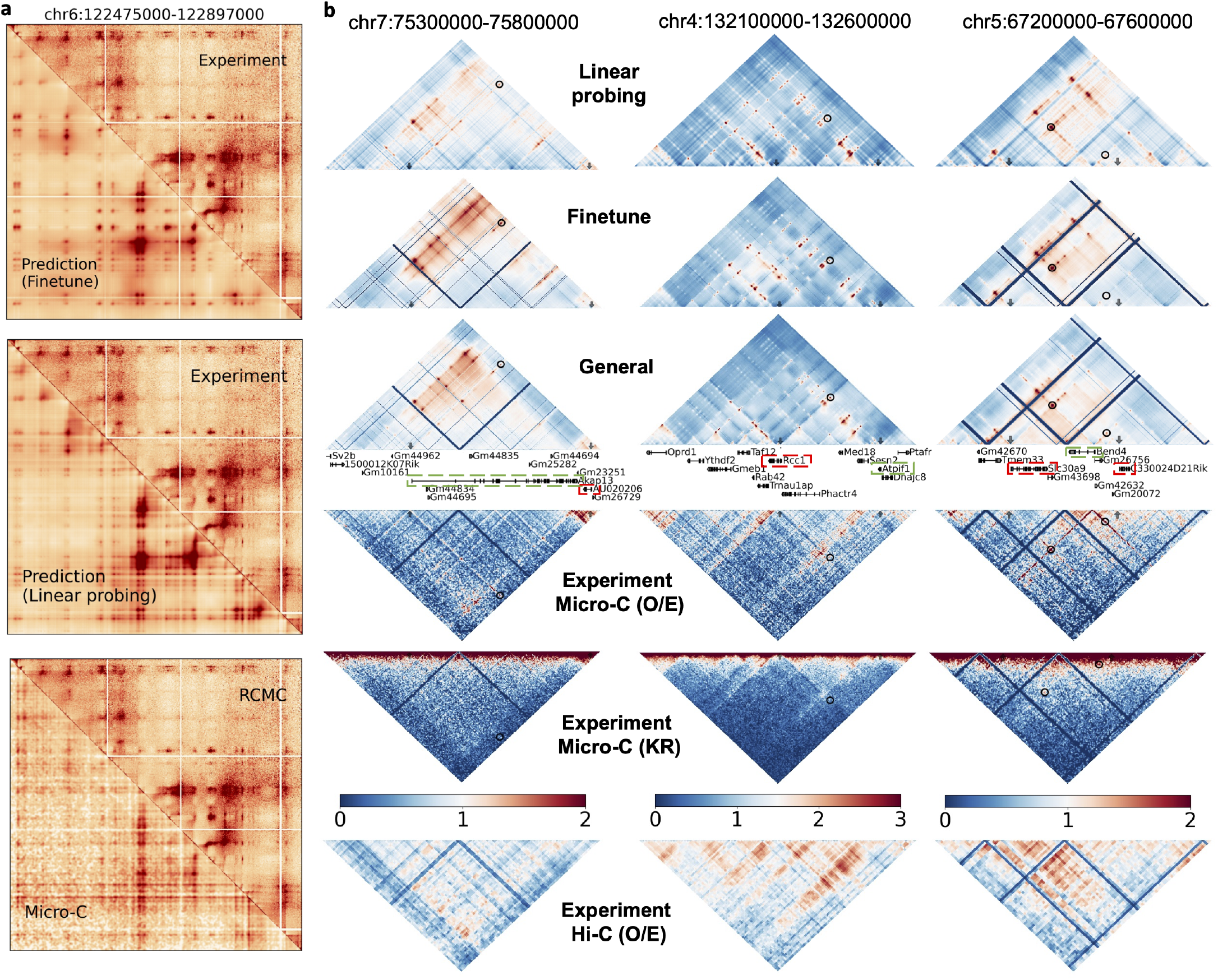
Evaluations on the predicted RCMC contact maps. **a**, Comparison of the RCMC experimental data with predictions from the fine-tuning model, linear probing model, and experimental Micro-C. **b**, Comparison of the predictions of four regulatory interactions from a promoter knockout study [55] among three approaches. The general model identifies all four interactions, while the fine-tuning model identifies three. Although the linear probing approach also identifies four interactions, the patterns are not as clear as those predicted by the general model. For example, the interaction between Bend4 and C330024D21Rik is less distinct. The KR-normalized Micro-C only clearly identifies one interaction. Although the interactions have a large signal in O/E-normalized Micro-C, they do not all exhibit loop patterns, such as in the BEND4 interaction example. Furthermore, the Hi-C contact map fails to detect all these interactions.

**Fig. S17.**
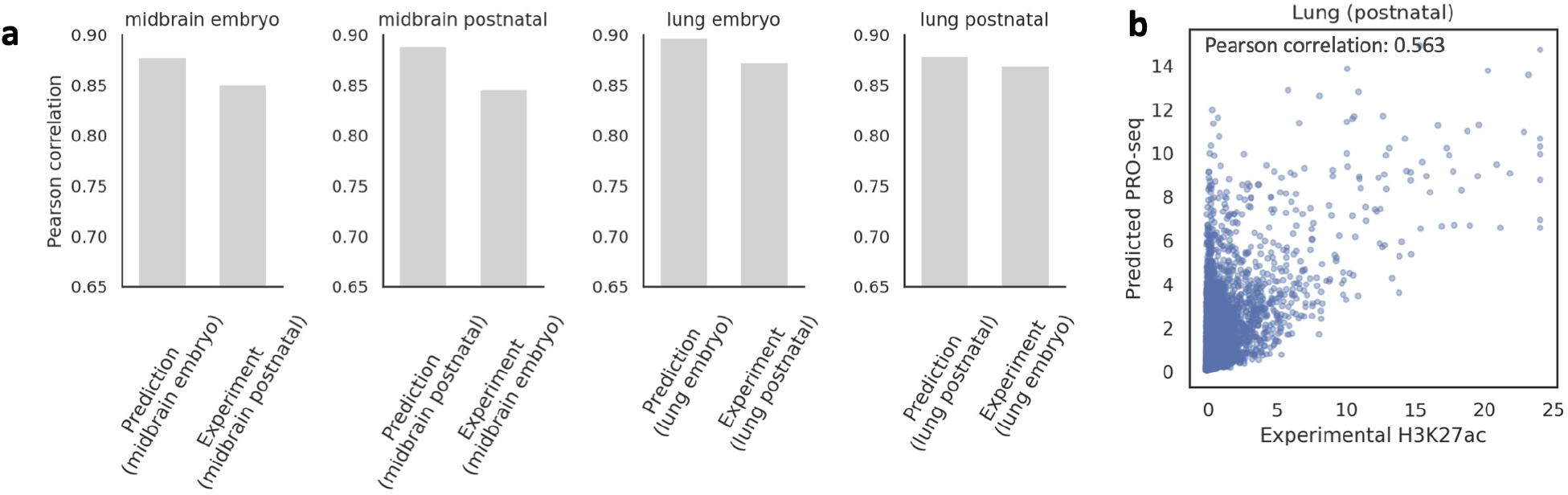
Evaluation of predicted gene expression and enhancer RNA signals. **a**, Pearson correlation scores comparing predicted and experimental RNA-seq data for midbrain and lung tissues across embryonic and postnatal stages. For each developmental stage (e.g., embryo), predictions based on experimental data from the same stage show higher correlations compared to using experimental data from a different stage. **b**, Correlation between predicted PRO-seq signals and experimental H3K27ac data across candidate enhancer regions in chromosome 1 of postnatal lung tissue.

**Fig. S18.**
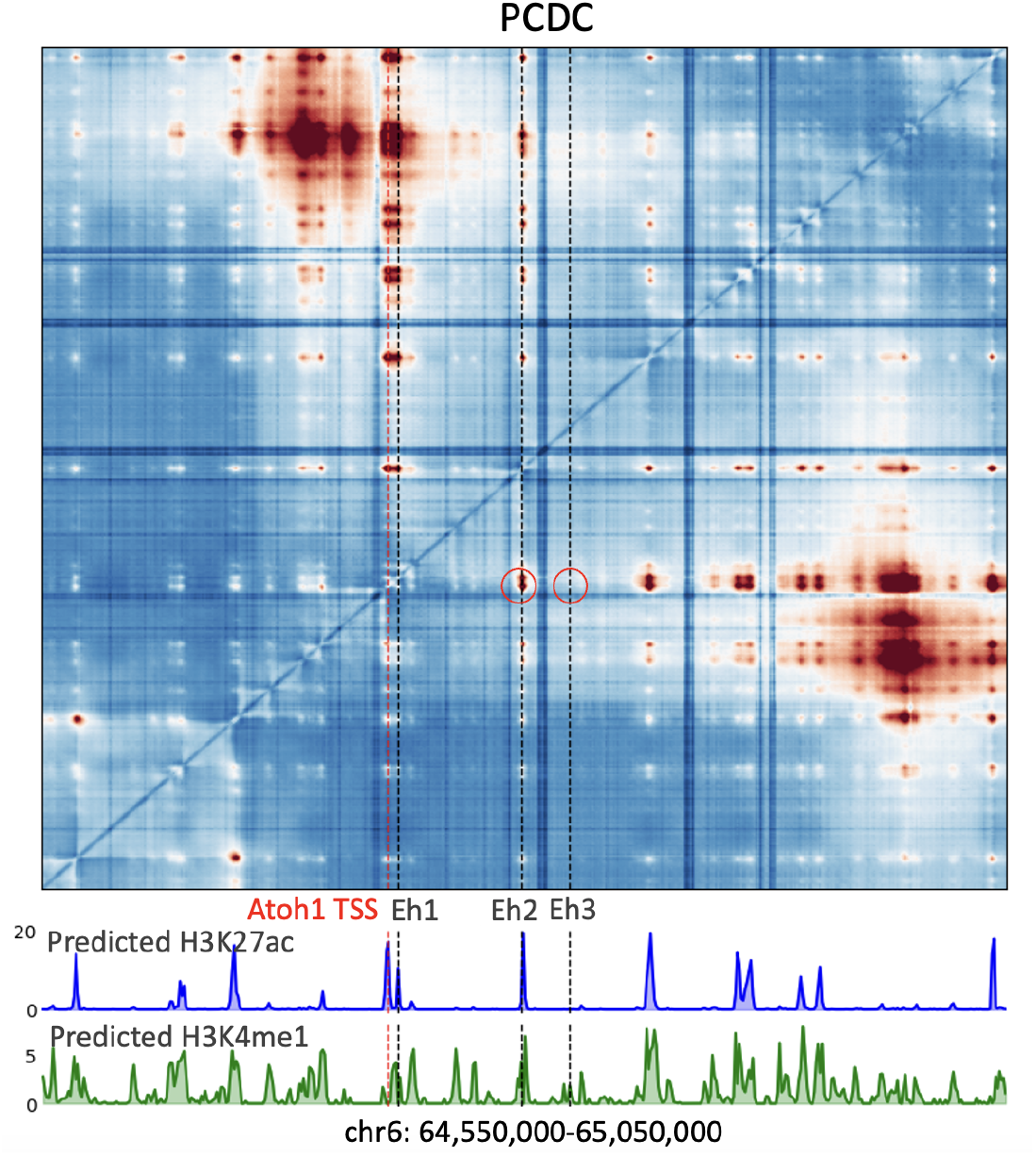
The model predicts a silencer at the En3 locus in the PCDC cell type. This locus lacks a clear predicted loop with the Atoh1 TSS and shows no predicted H3K27ac or H3K4me1 peaks, consistent with its silencer function.

**Table S1.**
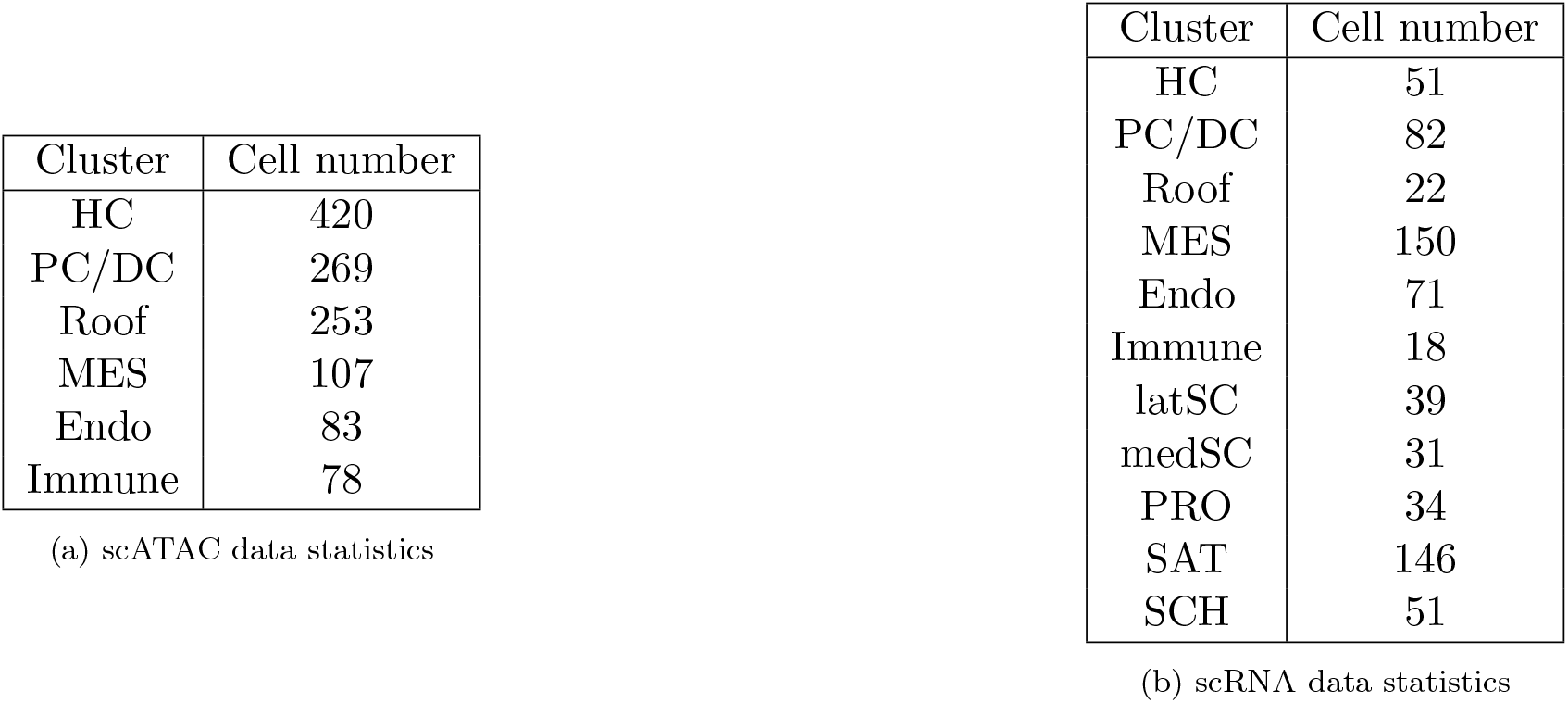
Statistics of scATAC data and scRNA data.

**Table S2.**
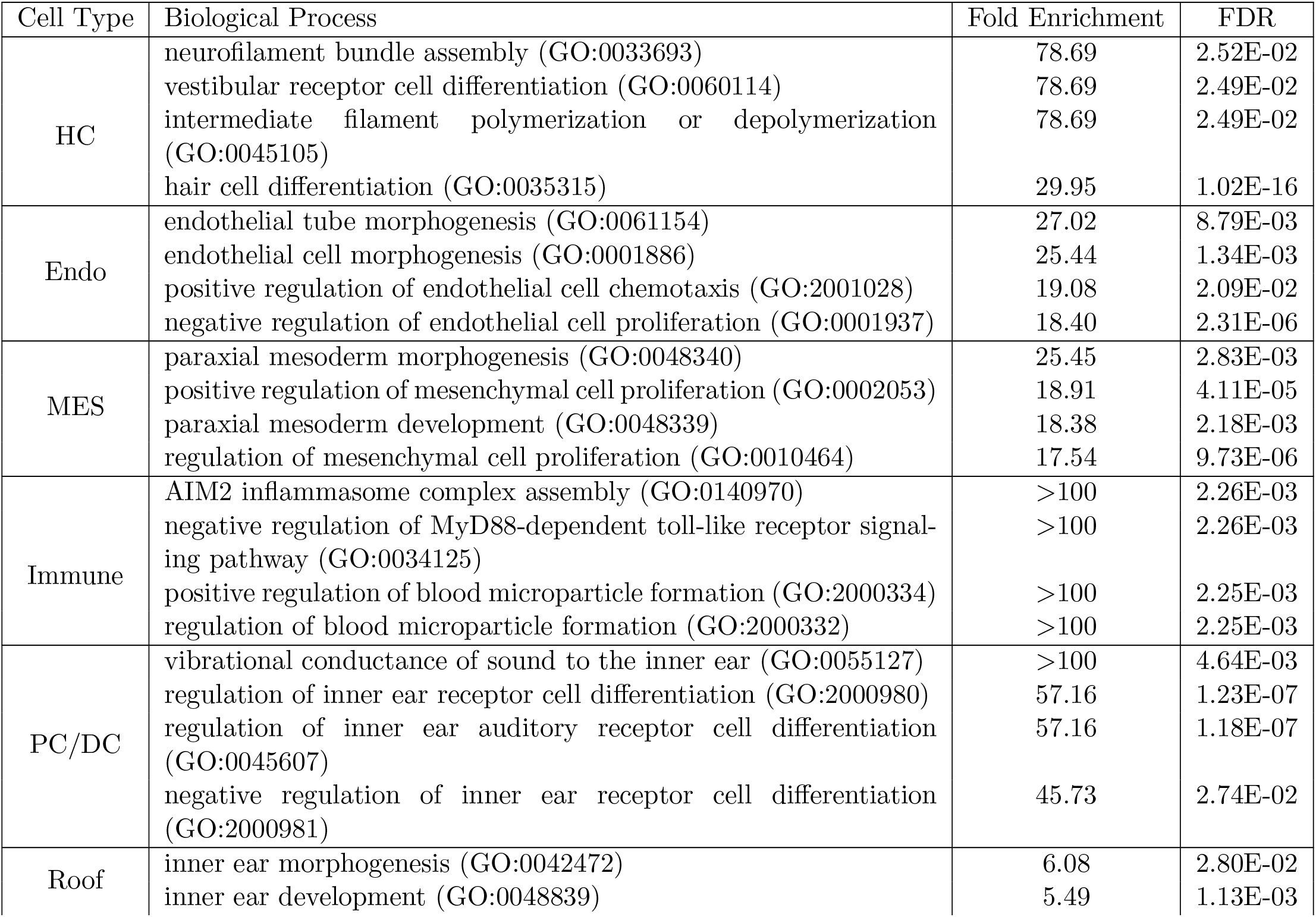
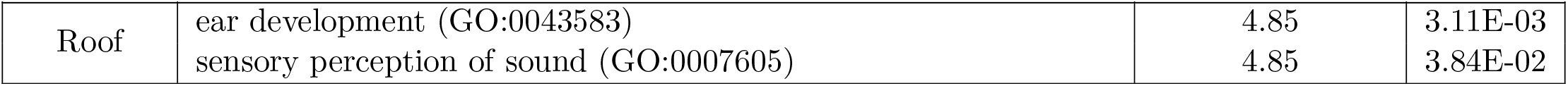
Inner ear GO biological process analysis shew the predicted gene expressions were accurate. In the HC cell type, processes such as ‘neurofilament bundle assembly’ and ‘vestibular receptor cell differentiation’ exhibit high fold enrichment and low false discovery rate, suggesting that these processes are highly characteristic of hair cells. Similarly, for the Endo cell type, processes like “endothelial tube morphogenesis” and “endothelial cell morphogenesis” are prominently enriched, aligning with the known functions of endothelial cells. The MES cell type shows enrichment in mesoderm development and mesenchymal cell proliferation processes, while the Immune cell type is associated with pathways related to inflammasome assembly and toll-like receptor signaling. Additionally, the PC/DC and Roof cell types display significant enrichment in processes involved in inner ear receptor cell differentiation and ear development, respectively. These enrichment results provide strong evidence that the selected marker genes effectively capture the distinct biological characteristics of each cell type, thereby validating the accuracy of the model’s predictions from a biological perspective.

| Cell Type | Biological Process | Fold Enrichment | FDR |
| --- | --- | --- | --- |
| HC | neurofilament bundle assembly (GO:0033693) | 78.69 | 2.52E-02 |
|  | vestibular receptor cell differentiation (GO:0060114) | 78.69 | 2.49E-02 |
|  | intermediate filament polymerization or depolymerization (GO:0045105) | 78.69 | 2.49E-02 |
|  | hair cell differentiation (GO:0035315) | 29.95 | 1.02E-16 |
| Endo | endothelial tube morphogenesis (GO:0061154) | 27.02 | 8.79E-03 |
|  | endothelial cell morphogenesis (GO:0001886) | 25.44 | 1.34E-03 |
|  | positive regulation of endothelial cell chemotaxis (GO:2001028) | 19.08 | 2.09E-02 |
|  | negative regulation of endothelial cell proliferation (GO:0001937) | 18.40 | 2.31E-06 |
| MES | paraxial mesoderm morphogenesis (GO:0048340) | 25.45 | 2.83E-03 |
|  | positive regulation of mesenchymal cell proliferation (GO:0002053) | 18.91 | 4.11E-05 |
|  | paraxial mesoderm development (GO:0048339) | 18.38 | 2.18E-03 |
|  | regulation of mesenchymal cell proliferation (GO:0010464) | 17.54 | 9.73E-06 |
| Immune | AIM2 inflammasome complex assembly (GO:0140970) | >100 | 2.26E-03 |
|  | negative regulation of MyD88-dependent toll-like receptor signaling pathway (GO:0034125) | >100 | 2.26E-03 |
|  | positive regulation of blood microparticle formation (GO:2000334) | >100 | 2.25E-03 |
|  | regulation of blood microparticle formation (GO:2000332) | >100 | 2.25E-03 |
| PC/DC | vibrational conductance of sound to the inner ear (GO:0055127) | >100 | 4.64E-03 |
|  | regulation of inner ear receptor cell differentiation (GO:2000980) | 57.16 | 1.23E-07 |
|  | regulation of inner ear auditory receptor cell differentiation (GO:0045607) | 57.16 | 1.18E-07 |
|  | negative regulation of inner ear receptor cell differentiation (GO:2000981) | 45.73 | 2.74E-02 |
| Roof | inner ear morphogenesis (GO:0042472) | 6.08 | 2.80E-02 |
|  | inner ear development (GO:0048839) | 5.49 | 1.13E-03 |
| Roof | ear development (GO:0043583) | 4.85 | 3.11E-03 |
|  | sensory perception of sound (GO:0007605) | 4.85 | 3.84E-02 |

